# Exact calculation of the expected SFS in structured populations

**DOI:** 10.1101/2023.05.10.540112

**Authors:** Armando Arredondo, Josué Corujo, Camille Noûs, Simon Boitard, Lounès Chikhi, Olivier Mazet

**Affiliations:** Institut National des Sciences Appliquées, Institut de Mathématiques de Toulouse, Université de Toulouse, Toulouse, France; Laboratoire Cogitamus; CBGP, Université de Montpellier, CIRAD, INRAE, Institut Agro, IRD, Montpellier, France; Instituto Gulbenkian de Ciência, Rua da Quinta Grande, No. 6, P-2780-156 Oeiras, Portugal; Laboratoire Évolution & Diversité Biologique (EDB UMR 5174), CNRS, IRD, UPS, Université de Toulouse Midi-Pyrénées, Toulouse, France

**Keywords:** Site frequency spectrum, population structure, structured coalescent, Markov processes

## Abstract

The Site Frequency Spectrum (SFS), summary statistic of the distribution of derived allele frequencies in a sample of DNA sequences, provides information about genetic variation and can be used to make population inferences. The exact calculation of the expected SFS in panmictic population has been derived in the Markovian coalescent theory for decades, but its generalization to the structured coalescent is hampered by the almost exponential growth of the states space. We propose here a complete algorithmic procedure, from how to build a suitable state space and sort it, to how to take advantage of the sparsity of the rate matrix and to solve numerically the linear system by an iterative method. The simplest case of the symmetrical *n*-island is then processed to arrive at a ready-to-use demographic parameters inference framework.

## 1. Introduction

Genomic data are increasingly used to reconstruct the recent evolutionary history of species [22, 8, 1]. To that aim methods vary either in the demographic models assumed for statistical inference [22, 12, 1] or in the type of genetic or genomic data used [2, 22, 8, 1]. One very popular way of summarizing genetic data is the site frequency spectrum (SFS), a histogram of allele frequencies which can be easily computed from genomic observed or simulated data [30, 11, 24, 12, 8]. While the expected SFS can be analytically predicted for panmictic populations [11], it cannot be derived for structured models with the exception of several important studies [4, 30, 19]. Whereas [4] derived the joint SFS for more than two populations, the demographic model used ignored gene flow, as in the two-population model of [30]. The more recent study of [19] focused on the joint SFS of two samples obtained from two populations connected by gene flow. The authors noted however that for larger sample sizes (*n*_1_ + *n*_2_ *>* 20) their approach would quickly become computationally impractical.

In this article we focus on establishing a general framework for predicting the expected aggregate (or marginal) SFS under a large family of structured models with many populations. This work is a continuation of the research we started a decade ago on the genealogical properties of genes sampled under the structured coalescent of Herbots [16, 32]. This led us to develop the concept of IICR (inverse instantaneous coalescence rate) which can be used to infer changes in connectivity under structured models [23, 6, 1]. While this previous work benefits from the fact that the IICR can be estimated from real data [22], it is limited by the fact that it focuses on the information provided by one pair of haploid genomes. Having multiple samples potentially provides better or more information on the recent past of the sample, since in typical genealogical trees, most of the coalescence times are distributed close to tree tips. Here, we show that the Markovian framework of the structured coalescent can be generalized to capture the necessary information for computing the expected SFS arbitrary numerical accuracy. The approach of [19] mentioned above limited itself to moderate sample sizes due to the computational challenges of handling the state spaces of the associated Markov processes. These state spaces grow in size almost exponentially as a function of the number of samples, making it difficult to compute the joint SFS for sample sizes greater than ten haploids per population for two populations.

Under the approach of [19] and assuming more than two populations the state space would become so large that the computation of the SFS would only be possible for much smaller sample sizes. For example, fixing the sampling size to 20 haploids, we show that the state space of Kern & Hey’s isolation-with-migration model with *n*_1_ = *n*_2_ = 10 is more than 260 times larger than what can be achieved under a model with 2 islands and any pattern of migration and sampling configuration by focusing exclusively on the aggregate SFS (further reductions in the size of the state space are possible under models with more symmetrical patterns of migration).

More generally, we show that by not tracking the historical origin of the lineages (mainly used to compute the joint SFS of the populations), using appropriate model specializations that take advantage of inherent symmetries, and exploiting the sparsity patterns that appear in the rate matrix under certain state orderings, it is possible to quickly compute the expected SFS of samples of size up to *k* = 26 haploids (thirteen diploids) in structured models such as the symmetrical *n*-island model under any sampling scheme and any number of populations.

The approach is general in the sense that it allows for many potential types of structured scenarios. However, we exclusively showcase the *n*-island as a model specialization for the implementation and results, once again due to its mathematical simplicity. Throughout the manuscript we propose novel methods for approaching the questions of demographic inference in the context of population structure, and we provide pseudocode for each of the algorithms discussed. Additionally, the concrete implementation is freely available at github.com/arredondos/sisifs.

### 1.1. The SFS

We can model the process in which mutations occur in genomic sequences in a simple way by introducing the assumption that the probability for a mutation to occur in the same position twice is null. This assumption is justified in the presence of very long sequences with low mutation rates, and it is known as the infinite sites model of mutation, in which mutations always happen in new locations. Under this assumption, we can assign binary states to the positions along the genome. The two states, which can be coded as 1 and 0, can be interpreted as the position containing a mutation or not, respectively. We can thus represent a genomic sequence as a string of zeroes and ones. If two genes are observed to have different codes in a given position, we say that the position is a segregating or polymorphic site. We could distinguish a mutated position from a non-mutated one, for instance, by comparing with a sampled sequence from an outgroup species.

A convenient way of summarizing the genetic diversity of a sample under these assumptions is to simply inspect the values of the sampled sequences in the segregating sites, given that the sequences match exactly in all other sites. Consider the following example, where *k* = 6 sequences were sampled, and 8 segregating sites were observed among them.

The rows of Table 1 contain the sequence encoding for each of the segregating positions (columns, labeled SNP *j* for *single-nucleotide polymorphism* number *j*). For instance, we can see that 4 of the 8 polymorphisms were *singletons*, meaning that a mutation happened in exactly one of the sampled sequences. We can similarly track the number of sites that exhibited exactly *i* mutations, and compile this information in a vector *ξ* = (*ξ*_1_, …, *ξ*_*k−*1_), where *ξ*_*i*_ is the number of segregating sites at which there are exactly *i* copies of the mutant type. In the example of Table 1, *ξ* = (4, 2, 1, 0, 1). Under the assumptions of a Wright-Fisher model with constant population size and scaled mutation rate *θ*, it can be shown that:

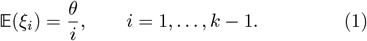

**Table 1:**
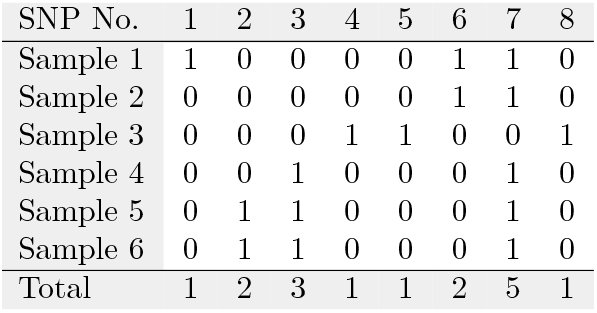
An example for how to compute the observed site frequency spectrum (SFS) in a sample of size *k* = 6 genes. The columns show the segregating sites (SNPs), which are the positions in the genome where a mutation is observed for at least one of the sampled sequences. In this case, the observed SFS is (4, 2, 1, 0, 1). With increased sequence length, the table grows horizontally, but the SFS is always of length k−1 because this is the maximum possible number of observed mutations (a column full of 1s is not possible since this would correspond to a mutation taking place before the most recent common ancestor of the sample, and as such it would not visible in the coalescence tree).

| SNP No. | 1 | 2 | 3 | 4 | 5 | 6 | 7 | 8 |
| --- | --- | --- | --- | --- | --- | --- | --- | --- |
| Sample 1 | 1 | 0 | 0 | 0 | 0 | 1 | 1 | 0 |
| Sample 2 | 0 | 0 | 0 | 0 | 0 | 1 | 1 | 0 |
| Sample 3 | 0 | 0 | 0 | 1 | 1 | 0 | 0 | 1 |
| Sample 4 | 0 | 0 | 1 | 0 | 0 | 0 | 1 | 0 |
| Sample 5 | 0 | 1 | 1 | 0 | 0 | 0 | 1 | 0 |
| Sample 6 | 0 | 1 | 1 | 0 | 0 | 0 | 1 | 0 |
| Total | 1 | 2 | 3 | 1 | 1 | 2 | 5 | 1 |

See Tajima [28], Fu [10], Griffiths and Tavaré [11], Hudson [18] for various proofs of (1). The vector *ξ* is known as the Site Frequency Spectrum, and it has been the basis for many estimators and test statistics for analyzing genomic data. Notable examples are the *θ*_*W*_ [31] and *θ*_*π*_ [27] estimators of *θ*; Tajima’s *D* [28]; Fay and Wu’s *H* [9]; and the fixation index *F*_*ST*_ [33].

### 1.2. Partitions

Counting the different ways a group of *k* lineages can be distributed among *j* islands is an important concept we use for enumerating the state space of our model. The theory of integer partitions is useful in this regard because it studies the ways an integer can be written as the sum of other integers.

We denote restricted partition numbers by *p*_*j*_(*k*). They count in how may ways can the number *k* be expressed as the sum of exactly *j* non-null numbers. In general, an integer partition of *k* is a multiset of positive integers that sum to *k*. For example, there are 7 possible partitions of *k* = 5, which can be represented as:

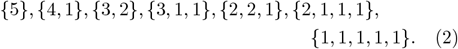

The order of the terms in the multiset is unspecified, so 5 = 2 + 3 is the same partition as 5 = 3 + 2. An alternative way of representing these partitions is by using *product notation*, where we specify the number of times each part is repeated, from 1 to *k* which is the largest possible part.

The partitions in (2) would be re-written as:

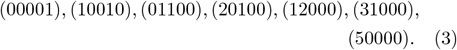

We denote by *P* (*k*), or simply *P* for simplicity, the total number of integer partitions of *k*. There is no known closed-form expression for *P*, but Hardy and Ramanujan [14] provided the asymptotic approximation 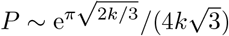. Algorithm P generates all partitions of *k* in standard notation (2). See Knuth [20, §7.2.1.4] for the derivation and detailed analysis of this algorithm.

#### Algorithm P

For evaluating routine `Partitions`(*k*), which visits, in reverse-lexicographic order, all integer partitions (*a*_1_, *a*_2_, …, *a*_*ω*_) such that 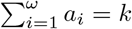.

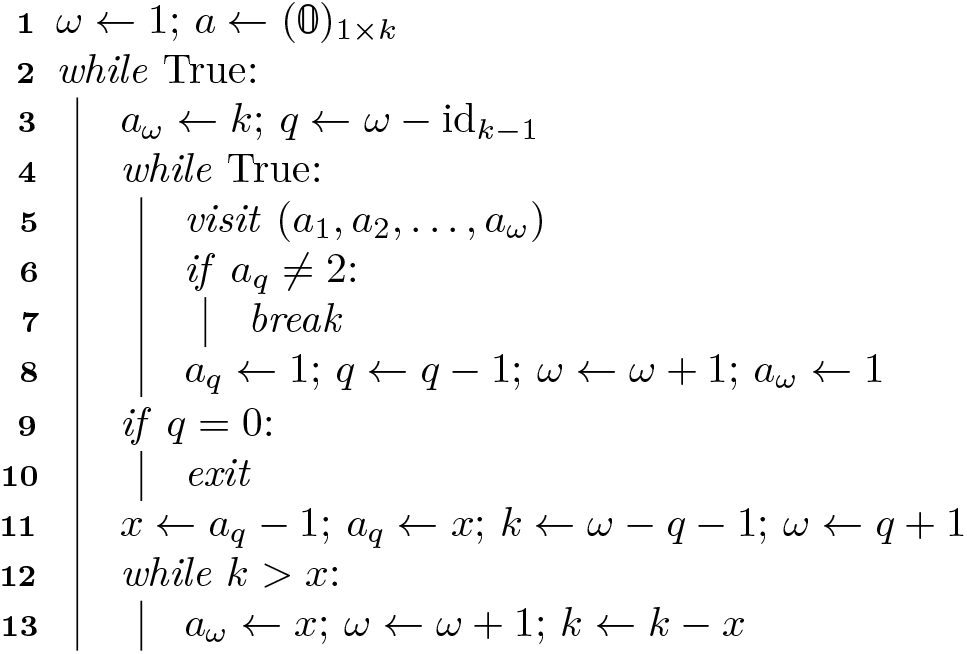

Integer partitions can be restricted in a multitude of ways. Here we are interested in restricting the number of parts. For instance, 5 = 2 + 3 is a partition in two parts, and 5 = 2 + 1 + 1 + 1 is a partition in four parts. The symbols *p*_*j*_(*k*) thus count the possible number of restricted partitions of *k* into *j* parts.

We may choose from these distinguishable or indistinguishable configurations depending on our needs. For instance, if the sizes of the islands are different in the general case, then the islands are inherently distinguishable. It is important to keep in mind that the choice of configuration will affect the size of the state space and the transition rates of the system. In any case, we denote the total number of states in the Markov chain by *E*.

## 2. The structured coalescent with ancestry tracking

Here we describe the state space for the continuous-time Markov process that models the migrations and ancestry of a sampled meta-population. We do this by introducing a generalization of the classical structured coalescent process.

The state space of the structured coalescent keeps track of what lineages are in which demes. For instance, an initial state of *α* = (0, 2, 1, 0) encodes that the model has *n* = 4 demes and *k* = 3 sampled lineages, with two of them located in deme 2 and the other in deme 3. We note that the islands are distinguishable by their index in the tuple, whereas the individual lineages are not (it is not possible to differentiate between the two lineages in deme 2 for instance).

This information is sufficient in order to compute the expected coalescence times 𝔼(*T*_*k*_), 𝔼(*T*_*k*−1_), …, 𝔼(*T*_2_) associated to the Markov process for any structured model. However, it is not enough information to compute a site frequency spectrum, since we would also need information about which lineages are ancestral to which throughout time. More precisely, in order to construct an SFS, we need to know at every instant how many descendants each lineage has in order to tell, given a mutation in an ancient lineage, how many copies it will have in the current sample. We achieve this by representing the states as matrices instead of tuples. We call these matrices *state matrices*.

### 2.1. State matrices

A state matrix has *k* rows and *n* columns, and the coefficient in row *i* and column *j* indicates how many lineages there are in deme *j* that are ancestral to *i* lineages from the sample. The example in Figure 1 shows one possible demographic history for *k* = 4 sampled lineages and *n* = 3 islands, with the corresponding state matrices.

**Figure 1:**
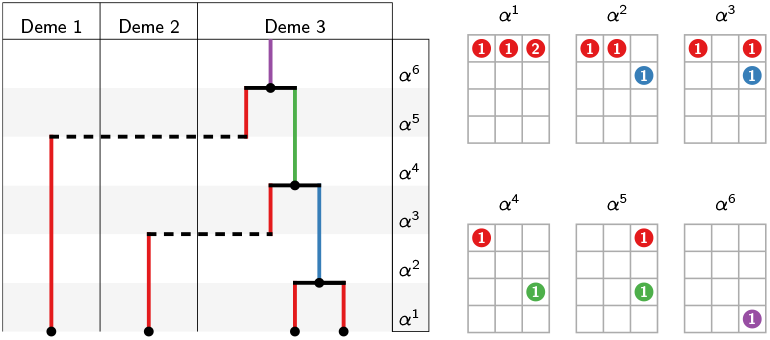
Example of a demographic history and the corresponding state matrices. In this case there are *n* = 3 demes in the model, and an initial sample of *k* = 4 lineages. Looking at state *α*^4^ for example, there is one lineage in deme 1 (column *j* = 1), which is ancestral to one lineage (row *i* = 1) from the sample. There is also one lineage in deme 3 (column *j* = 3), which is ancestral to three lineages (row *i* = 3) from the sample. Thus, any mutation to this (green) lineage in *α*^4^ will be present in three of the four sampled lineages, and will thus contribute to coefficient *ξ*_3_ of the SFS. The non-zero coefficients of the state matrices and their corresponding lineages in the coalescence tree have been color-coded in this figure according to the number of ancestral lineages or *weight* for ease of reference. The remaining null coefficients of the matrix are omitted in order to avoid visual clutter. The migration rates, deme sizes and coalescence times of the model are not represented.

Given our description of state matrix for a model with *n* demes and *k* samples, there is a relationship between the number of sampled lineages and the coefficients *α*_*ij*_ of any state matrix *α*, which is given by the invariant:

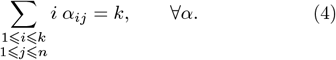

We refer to the factor *i* in (4) as *the weight* of the lineages that are in the row *i*. It represents the fact that each lineage in row *i* is ancestral to *i* lineages from the sample. The number of lineages that are present in any given state *α* is denoted |*α*|, so:

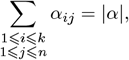

and we call *r*_*α*_ the number of coalescences that have taken place up to state *α*, such that *k* = |*α*| + *r*_*α*_ for all *α*. We note that the state matrix associated with any state *α* is empty for rows below *r*_*α*_ + 1. Indeed, if we suppose *α*_ij_⩾1 for some *i ⩾r*_*α*_ + 2, then the weight of that lineage is at least *r*_*α*_ + 2, and since the remaining |*α*| − 1 lineages have weight at least 1, then the total weight of the state is:

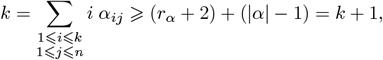

which is absurd.

We denote by *ε* the set of all possible states *α*, and by *E* the total number of states:

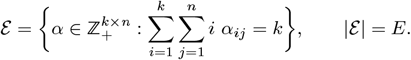

The set *ε* is clearly defined by the sample size *k* and number of demes *n*, but in order to avoid notation clutter we keep this dependency implicit.

Another important notion related to the state matrices are the contribution vectors. The contribution vector of state *α*, denoted *ψ*_*α*_, is defined as the sum by rows of the states matrices:

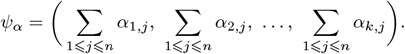

Returning to the example from Figure 1, the contribution vectors are:

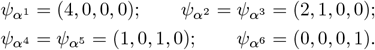

In general, for *i < k*, the coefficient *ψ*_*α*_(*i*) represents the contribution to 𝔼(*ξ*_*i*_) for each unit of time the process spends in state *α* on average (see §3). We denote by *𝒫* the set of all possible contribution vectors, and by *P* its total number of elements:

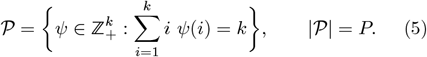

In this case, 𝒫 depends only on the number of samples *k* (in fact 𝒫 = *ε* when *n* = 1), but once again we keep this dependency implicit.

### 2.2. Transitions between states

From this point onwards we use the notation *e*_*ij*_ to indicate a matrix of size *k* × *n* with null coefficients except for the one in row *i* and column *j*, which has a value of 1. There are two possible types of state transitions: migration events and coalescence events. During a migration event, one lineage in deme *j* migrates to deme *g* ≠ *j* with rate *m*_*jg*_. We collect these rates in the matrix

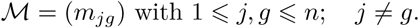

We convene, for the sake of completeness, that the diagonal elements of ℳ are null.

Given two states *α* and *β*, and a weight of *i ∈* {1, …, *k*− 1}, the process can transition from *α* to *β* via migration event if for some *j, g ∈* {1, …, *n*}

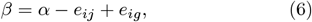

and this transition happens with rate *m*_*jg*_ for every lineage that could potentially migrate, of which there are *α*_*ij*_. Algorithm M generates all possible destination states *β* given any departure state *α*.

#### Algorithm M

For evaluating routine `Migration Destinations`(*ℳ, α*), which given the migration rates and a state matrix *α* of size *k* × *n*, visits all the tuples (*β, m*) where *β* is a state satisfying (6) and *m* (which may be null) is the corresponding transition rate.

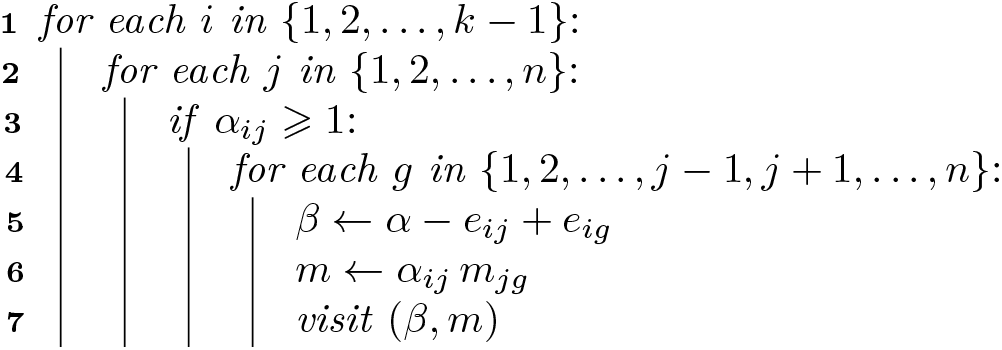

Algorithm M visits all *potential* migration destinations, regardless of whether a transition would be possible given the migration rates ℳ. This behaviour is desired because, in addition to generating the migration rates, we use the algorithm to build the state space *ε* by repeatedly simulating all possible migrations (see §2.4 and Algorithm A).

A coalescence happens when two lineages in the same deme *j* are replaced with their immediate common ancestor, also in deme *j*. This coalescence happens with a rate that is the inverse of the deme size. We collect the deme sizes in the vector

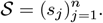

Given two states *α* and *β*, and two values *i, h ∈* {1, …, *k* − 1}, the process can transition from *α* to *β* via coalescence event if for some *j ∈* {1, …, *n*}

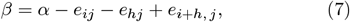

and this transition happens with rate 1*/s*_*j*_ for every pair of lineages that can coalesce, of which there are 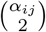 if *h* = *i*, and *α*_*ij*_ *α*_*hj*_ otherwise. Algorithm C generates all possible states to which the process can transition by a coalescence from *α*.

#### Algorithm C

For evaluating routine `Coalescence Destinations`(*𝒮, α*), which given the deme sizes and a state matrix *α*, visits all the tuples (*β, c*) where *β* is a state satisfying (7) and *c* ≠ 0 is the corresponding coalescence rate.

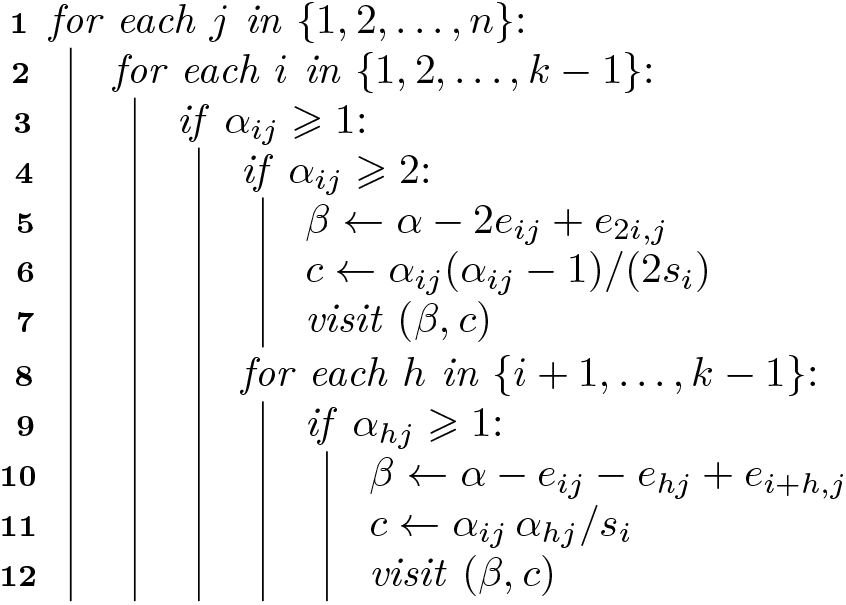

### 2.3. Grouping and sorting the states

We now focus on defining the rate matrix of the Markov process. This is a matrix *Q* of size *E* × *E*, where coefficient (*ℓ*_0_, *ℓ*_1_) contains the rate of transition from state 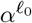 to state 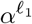 when *ℓ*_0_ ≠ *ℓ*_1_ and the diagonal elements are defined such that 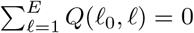. To define such a matrix, we first need to define an order ≺ on the set of all states *ε*. The focus of this section will therefore be the construction of a bijective map

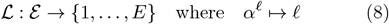

such that 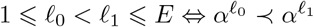.

We start with the partial order given by the natural transition of the process from states with more living lineages to states with fewer living lineages. Indeed, migration events do not change the number of living lineages, and a coalescence events decrease them by one, therefore |*α*| (or equivalently, *r*_*α*_) decreases (increases) during any given instantiation of the process. We convene then that:

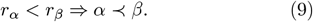

In the example from Figure 1, since 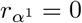 and 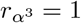, we have *α*^1^ ≺ *α*^3^.

By grouping together the states that share the same number of live lineages, we get a partition of the state space into *k* disjoint parts (one part for each possible value of *r*_*α*_ from 0 to *k* − 1). We denote these parts by *ε*_*i*_, each with *E*_*i*_ elements, such that *ε* = ∪_*i*_ *ε*_*i*_ and *E* = Σ_*i*_ *E*_*i*_:

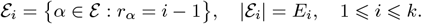

The partition {*ε*_*i*_} can be understood as the quotient set *ε/R*, where *R* is the equivalence relation

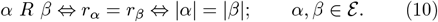

This partial order imposes a *k* × *k* block structure on *Q*, where block (*i, h*) has dimensions *E*_*i*_ × *E*_*h*_, and contains the rates of transitioning from the states in *ε*_*i*_ to the states in *ε*_*h*_.

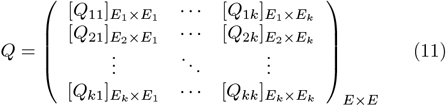

We note that if the process is at a non-absorbing state *α* ∈ *ε*_*i*_, then it can only transition to another state in *ε*_*i*_ via migration, or to a state in *ε*_*i*+1_ via coalescence, therefore all the coefficients of block [*Q*_*ih*_] are null when *h* ≠ *i* and *h* ≠ *i* + 1. Matrix *Q* is thus block bi-diagonal, with blocks *A*_*i*_ = [*Q*_*ii*_] for 1 ⩽ *i* ⩽ *k* in the diagonal, and blocks *B*_*i*_ = [*Q*_*i, i*+1_] for 1 ⩽ *i* ⩽ *k* − 1 in the upper diagonal.

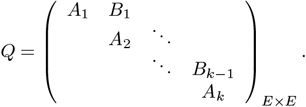

We call *A*_*i*_ the migration blocks and *B*_*i*_ the coalescence blocks of *Q*. We further note that migration block *A*_*k*_ is null, since there are no more transitions once the process arrives at any of the stationary states of *ε*_*k*_. An alternative representation of *Q* is thus possible:

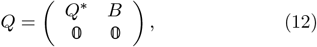

where

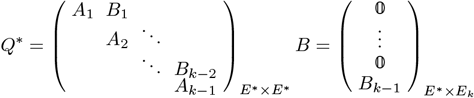

Here, *Q*^*\**^ is known as the sub-intensity matrix and contains the rates of transition between the transient states *ε*^*\**^ = *ε* − *ε*_*k*_. *E*^*\**^ is the number of transient states *E* − *E*_*k*_. Next we focus on defining the order ≺ for the states within the sets *ε*_*i*_. We begin by defining another equivalence relation *C* between states of *ε*. We say that two states are equivalent under *C* if they have the same contribution vector:

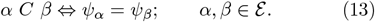

This relation partitions the state space into disjoint sets identified by their common contribution vector. We can thus think of each different contribution vector as the representative of an equivalence class of states, or a *macrostate*. In order to count how many different macrostates there are in a process with sample size *k*, we need to briefly return to integer partitions. It is clear from their product representation (3) that the set of all integer partitions of *k* is the set 𝒫 of all possible state contribution vectors defined in (5). There are thus *P* possible macrostates in the process. The number of parts in macrostate *ψ*_*α*_ ∈ *𝒫* is given by 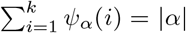. We group the macrostates by their number of parts following a similar procedure to how we group the states by their number of live lineages in (10). Overloading the name *R* to define a relation on 𝒫, we say that:

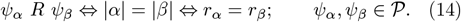

This partitions 𝒫 into *k* disjoint sets 𝒫_*i*_, each with *P*_*i*_ macrostates, such that 𝒫 = ∪_*i*_ 𝒫_*i*_ and *P* = Σ_*i*_ *P*_*i*_:

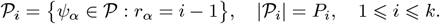

We sort the macrostates of 𝒫_*i*_ in reverse lexicographic order of standard notation (2) as does Algorithm P, so for *k* = 5 we have (4, 1) ≺ (3, 2) in 𝒫_2_, and (3, 1, 1) ≺ (2, 2, 1) in 𝒫_3_. This is a total order in 𝒫_*i*_, so for each *i* ∈ 1, …, *k* we will assume the existence of a bijective map ℒ_*i*_ defined on 𝒫_*i*_ such that ℒ_*i*_(*ψ*) ∈ 1, …, *P*_*i*_, and for *ψ, φ* ∈ 𝒫_*i*_, we have *ψ* ≺ *φ* ⇔ *ℒ*_*i*_(*ψ*) *< ℒ*_*i*_(*φ*).

Since *ψ*_*α*_ *∈ ℒ*_*i*_ ⇔ *r*_*α*_ = *i* − 1 ⇔*α* ∈ *ε*_*i*_, this order in 𝒫_*i*_ motivates a natural way to introduce a partial order in the elements of *ε*_*i*_. We convene that for *α, β* ∈ *ε*_*i*_,

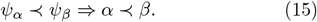

We can group together the uncomparable states of *ε*_*i*_ by returning to the equivalence relation *C* defined in (13) such that *α C β ⇔ ψ*_*α*_ = *ψ*_*β*_ for *α, β* ∈ *ε*_*i*_. There are *P*_*i*_ equivalence classes in the quotient set *ε*_*i*_*/C*, which we will denote *ε*_*ip*_, each with *E*_*ip*_ states. We now have 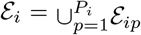 and 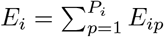.

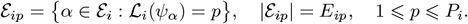

As was the case with *Q* in (11), this partial order on *ε*_*i*_ imposes a *P*_*i*_ × *P*_*i*_ block structure on migration block *A*_*i*_, and a *P*_*i*_ × *P*_*i*+1_ block structure on coalescence block *B*_*i*_. For *p, q* ⩽ *P*_*i*_, sub-block (*p, q*) of *A*_*i*_ has dimensions *E*_*ip*_ × *E*_*iq*_, and for *q*^*′*^ ⩽ *P*_*i*+1_, sub-block (*p, q*^*′*^) of *B*_*i*_ has dimensions *E*_*ip*_ × *E*_*i*+1, *q ′*._

Upon closer inspection, we see that the contribution vectors of the states in *ε*_*i*_ are invariant under migration. Indeed, if *β* = *α − e*_*ij*_ + *e*_*ig*_, then the sum of row *i* of both states remains unchanged, and thus *ψ*_*α*_ = *ψ*_*β*_. This implies that sub-block (*p, q*) of migration block *A*_*i*_ has all null entries except when *p* = *q*, and thus *A*_*i*_ is *P*_*i*_ *× P*_*i*_ block-diagonal. We denote the diagonal sub-blocks of *A*_*i*_ as 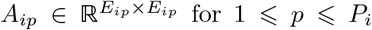 for 1 ⩽ *p* ⩽ *P*_*i*_. There is no such simplification for coalescence block *B*_*i*_, but we group all its sub-blocks in each block-column as 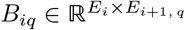 with 1 ⩽ *q* ⩽ *P*_*i*+1_ for later convenience:

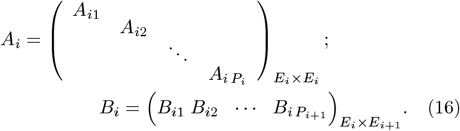

To make ≺ a total order, we need to define it for the states within each of the *P* sub-sub-spaces *ε*_*ip*_. There is no further benefit in doing so in terms of the structure of the rate matrix *Q* in the general case, but it is still a necessary step for building it. We thus convene that the order of the states in *ε*_*ip*_ is the one obtained by applying algorithm M to state 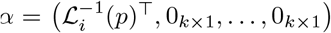 and all its subsequent migration destinations repeatedly (this procedure is detailed in algorithm A). Therefore, for every *i* = 1, …, *k* and *p* = 1, …, *P*_*i*_ we assume the existence of a bijective map *ℒ*_*ip*_ defined on *ε*_*ip*_ such that *ℒ*_*ip*_(*α*) ∈ {1, …, *E*_*ip*_} and *α ≺ β* ⇔ *ℒ*_*ip*_(*α*) *< ℒ*_*ip*_(*β*). With this construction, we can finally map every state *α ∈ ε* to an index *𝓁 ∈* {1, …, *E*}, and the definition of the global map *ℒ* defined in (8) would be

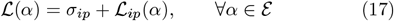

where:

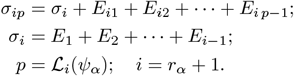

Figure 2 presents a schematic summary of some of the objects introduced in this section, their relations and the associated notation.

**Figure 2:**
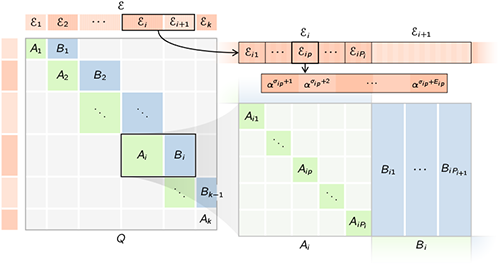
Schematic representation of the macro- and micro-structure of the state space *ε* and the rate matrix *Q* that results from appropriately grouping and sorting the states.

### 2.4. Construction of the rate matrix

We now propose a method for the construction of the migrations sub-blocks *A*_*ip*_, coalescence sub-blocks *B*_*ip*_ and indexing sub-maps *ℒ*_*ip*_ by combining the algorithms of the previous section. The method is structured in two separate stages.

#### Discovery and migration stage

The goal of this stage is to generate all the states in a constructive manner by following all possible migration paths, collecting at the same time the information of the transition rates in the migration sub-blocks *A*_*ip*_.

We know from the previous section that the state space is partitioned by the relation *C* from (13) into *P* sub-subspaces denoted *ε*_*ip*_, with no possible migration between them. We thus proceed to *discover* the states by initializing the sets *ε*_*ip*_ with the class representatives of *ε /C*. These are the states that have all their live lineages in the first deme, so they can always be defined regardless of the value of *n* by placing their associated contribution vector in the first column:

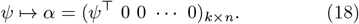

The contribution vectors *ψ* are obtained by converting the partitions (*a*_1_, …, *a*_*ω*_) generated by algorithm P to product notation (3). See lines [3 … 6] in Algorithm Q.

The next step is to generate (and add to *ε*_*ip*_) all the states that can be reached via migration from the initial state (18) using Algorithm M, then all the states that can be reached from those new states, and so on until no new states are being visited or added. This operation is accomplished using a *queue* data structure, or any generic data structure that implements the FIFO (first in, first out) interface, where elements are added at one end of the queue, and removed from the other end.

For our use, we may represent a queue of states as a tuple of length 0 ⩽ *ω* < ∞ with elements in *ε*, and two associated functions: enq and deq, which add (enqueue) and remove (dequeue) elements from the queue, respectively. More precisely, we say that *U* ∈ *ε*^*ω*^ is a queue of size *ω*; and for 𝒰 = *⋃*_*ω⩾*0_ *ε*^*ω*^ we define the maps

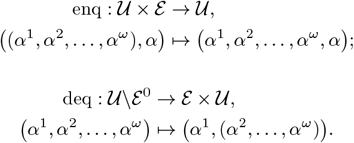

For a formal treatment of the queue abstract data structure (and other common data structures in computer science), see Lehmann and Smyth [21].

Each time a state *β* is found to be reachable via migration from state *α*, we accumulate the corresponding migration rate *m* in coefficient *A*_*ip*_(ℒ_*ip*_(*α*), *ℒ*_*ip*_(*β*)) of the migration sub-block. The index mapping functions ℒ_*ip*_ are also progressively constructed. We represent them as subsets of the cartesian product *ε*_*ip*_ × {1, …, *E*_*ip*_}, which contains tuple (*α, 𝓁*) if and only if *ℒ*_*ip*_(*α*) = *𝓁*. Building these data structures is sufficient to fully define the state space, since the state spaces *ε*_*ip*_ are simply the domains of the maps, which we reference using the notation 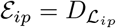. Algorithm A summarizes the proposed procedure for this stage.

##### Algorithm A

Defines `MigrationSubBlock`(*ℳ, α*^0^), which given the pairwise migration rates *ℳ* of the island model and a protostate *α*^0^ (18) in *ε*_*ip*_, builds and returns (*A*_*ip*_, *ℒ*_*ip*_).

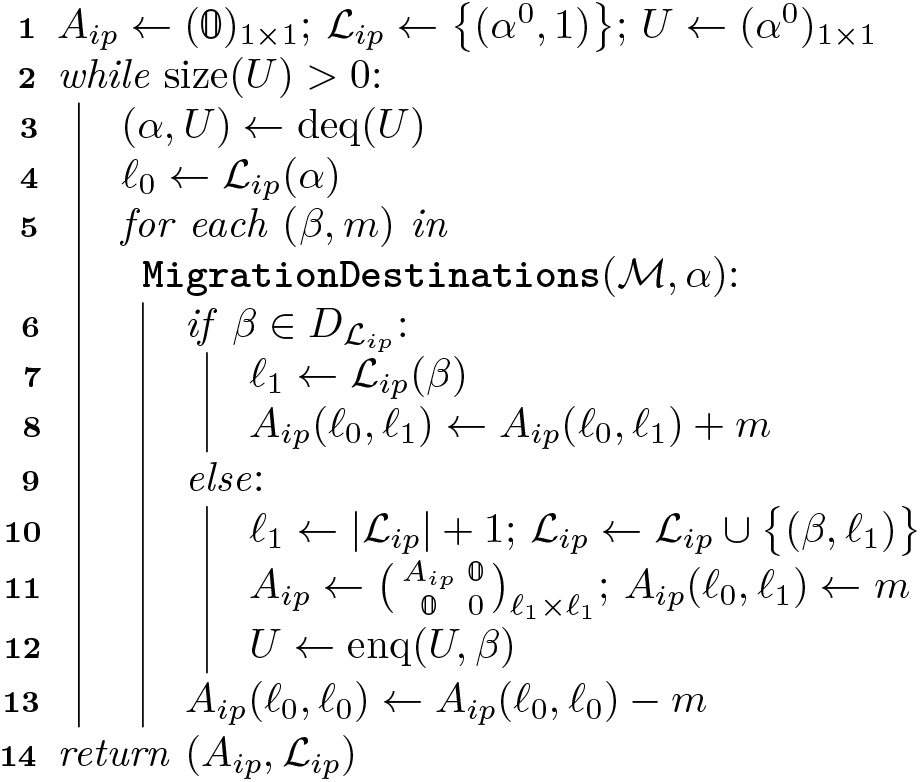

#### Coalescence stage

The goal for this stage is to compute the coalescence sub-blocks *B*_*ip*_, and consequently update the diagonal entries of sub-blocks *A*_*ip*_.

Since at this point all the states *ε*_*ip*_ are known, as well as the index maps ℒ_*i*_ and ℒ_*ip*_, the procedure simply consists in visiting, for every state *α*, all possible states *β* that can be reached by coalescence. This is accomplished by using Algorithm C and accumulating the coalescence rates in coefficient *B*_*iq*_(*σ*_*ip*_ − *σ*_*i*_ + ℒ_*ip*_(*α*), *ℒ*_*i*+1, *q*_(*β*)) for the corresponding value of *q*. The detailed procedures are presented in lines [9…18] of Algorithm Q. In general, Algorithm Q presents the complete construction of the Markov process, showing the details of how to combine all algorithms specified so far. Here, we omit some of the more menial details of the process, which includes how to compute the sizes *E*_*i*_ and *E*_*ip*_ given the existing information, and how to assemble the final rate matrix *Q* and index map *L* by arranging their respective sub-parts *A*_*ip*_, *B*_*ip*_ and ℒ_*ip*_.

## 3. Computing the expected SFS

In this section we show, given the Markov process defined previously, how to compute the expected SFS for an arbitrary model structure and sample configuration. This computation is reduced to a linear algebra problem for which we propose an efficient solution in Algorithm X.

### Algorithm Q

Defines `RateMatrix`(*k, n*, ℳ, 𝒮), that given the model parameters, builds the rate matrix of the Markov process and the associated data structures.

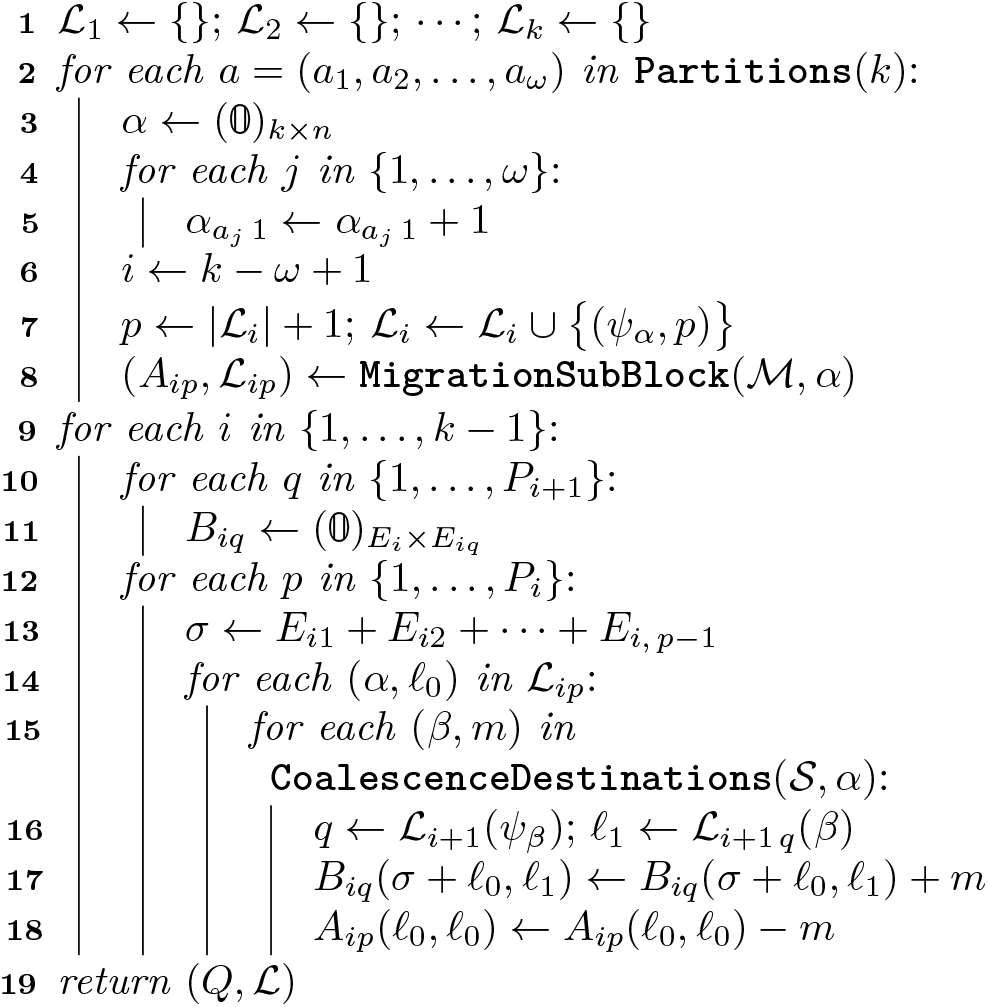

In the infinite-sites model, all mutations occur on unique sites, and as such the number of mutations is equal to the number of segregating sites. When a mutation occurs in a lineage with *i* descendants in the sample (a lineage of weight *i*), the corresponding segregating site has *i* copies of the mutated allele. Additionally, since we model mutations as a Poisson process of intensity *θ* along each branch of the genealogy, the total number of mutations that occur in lineages of weight *i* has an expected value of *θ* times the expected total length of such lineages, which we denote as *ζ*_*i*_:

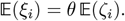

In the example of Figure 1, *ζ*_1_, *ζ*_2_ and *ζ*_3_ would be the total length of red, blue and green branches, respectively.

Since we are interested in the normalized SFS 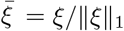, the parameter *θ* is cancelled away, so we focus exclusively on the quantities 𝔼(*ζ*_*i*_). These can be computed by adding, for every transient state *α* ∈ *ε*\ *ε*_*k*_ = *ε* *, the number of lineages of weight *i* (given by *ψ*_*α*_(*i*)) multiplied by the expected total time spent in the state before absorption, which we denote by *T*_*α*_. We thus have:

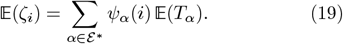

In order to compute the times 𝔼(*T*_*α*_), we return to the sub-intensity matrix *Q*^*^ defined in (12). This matrix is nonsingular, and *G* = (−*Q*^*^)^−1^, known as the process’ Green matrix, has the property that coefficient *G*(*ℓ*_0_, *ℓ*_1_) is the expected time that the process spends in state 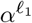 before absorption, given an initialization at state 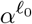 (see Bladt and Nielsen [3], Theorem 3.1.14). With an initial distribution for the transient states *π*^0^ = *π*^0^(1), *π*^0^(2), …, *π*^0^(*E*^*^)^⊤^, we can then write 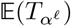 as

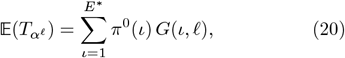

where *E*^*^ = |*ε*\*| = *E* −*E*_*k*_. We can combine and rearrange equations (19) and (20) to get

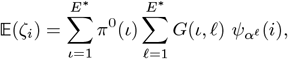

which can be further simplified when considering all values of 1 ⩽ *i < k* as

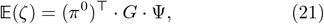

where Ψ, called the contribution matrix, is an *E*^*^ *×* (*k* −1) matrix where coefficient 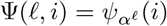 is the number of lineages of weight *i* in state *α*^*ℓ*^.

### 3.1. Numerical solution

We now turn to the question of how to numerically solve problem (21). The computation of matrix *G* involves inverting *Q*^*^, which grows exponentially in size with the number of the samples *k* (see Figure 4), so it is inevitable that the computational cost of the solution will eventually exceed the available resources regardless of the underlying method, given a large enough value of *k* = *k*_critical_. Due to this inherent and unavoidable complexity, our strategy simply consists in exploiting the sparsity and structure of *Q* (§2.3) to delay *k*_critical_ as much as possible.

We avoid the direct computation of the inverse *G*. Instead, we compute the product (*π*^0^)^⊤^ ·*G* by solving the linear system

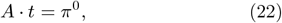

where *A* = (−*Q*^*^)^⊤^. The unknowns of this system are the coefficients of a vector *t*∈ R^*E*\* × 1^, which contains in component *ℓ* the time 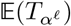 given initial distribution *π*^0^. Following the notation conventions of §2.3, we denote by *t*_*i*_ and 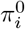 the segments of vectors *t* and *π*^0^ corresponding to states in *ε*_*i*_, for *i* = 1, …, *k* − 1; and by and 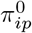 the subsegments corresponding to the states in *ε*_*ip*_ for *p* = 1, …, *P*_*i*_. We can thus write system (22) in the following block form:

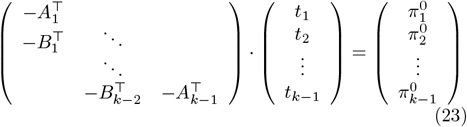

This structure suggests the sequence of sub-problems:

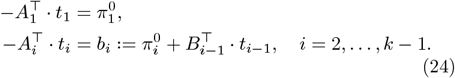

We note that in (24), the problems must be solved sequentially, since the solution segment *t*_*i*_ depends on the previous one *t*_*i* −1_. Assuming thus that *i ⩾*2 and *t*_*i*− 1_ is known, we now focus on computing *t*_*i*_. Exploiting the sub-structure of the system matrix *A*_*i*_ (16), the problem 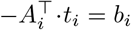 can be decomposed into the following *P*_*i*_ sub-problems (Figure 2):

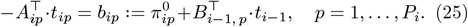

We note that the computation of the solution sub-segment *t*_*ip*_ does not depend on any other sub-segments of *t*_*i*_; only on the already-available segment *t*_*i*−1_. This implies that all *P*_*i*_ problems in (25) can be solved in parallel. For each one of them we use the Successive Over-Relaxation iterative method: given an initial guess 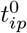, the scheme (known as SOR(*ω*)) generates a sequence of solutions 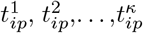, where coefficient 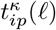 is computed using information from 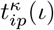 for *ι < ℓ* and 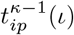 for *ι > ℓ*, so there is no need to keep two versions of the solution sub-segment *t*_*ip*_ in memory. We use the norm of the residual vector 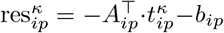 as the convergence metric, so for a given desired precision *δ*, the stopping criterion is 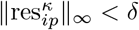. We denote by 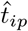 the final approximation 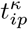 of the unknown vector *t*_*ip*_. The detailed procedure is given in lines [7 ··· 10] of Algorithm X.

The parameter *ω* in the SOR method affects the rate of convergence. A necessary condition for convergence is 0 *< ω <* 2, and for system matrices that are *M* -matrices (as is the case for all the *A*_*ip*_), a sufficient condition for convergence is 0 *< ω* ⩽ 1 (see §7.2.3 in Hackbusch [13]). In practice, values of *ω* larger than 1 can be used to accelerate convergence despite not satisfying the sufficient convergence condition.

Algorithm X summarizes our proposed method for computing the expected SFS 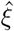 given the rate matrix *Q* of the Markov process and an initial state distribution *π*^0^.

### 3.2. Error analysis

Algorithm X guarantees that after each final iteration of the SOR scheme, the residues 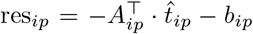 satisfy ∥res_*ip*_ ∥_∞_*< δ*. It is then easy to verify that the global residue of system (22) given by 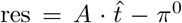 satisfies ∥res∥_∞_ *< δ*. This is a consequence of the fact that the vectors res_*ip*_ are simply sub-segments of res, and that for *X*⊂ℝ we have max_*x*∈ *X*_ |*x*| *< δ* ⇔ ∀*x* ∈ *X*, |*x*| *< δ*. We can thus say that given *δ >* 0, Algorithm X computes 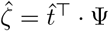, an approximation of 𝔼(*ζ*) (21) where

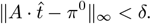

#### Algorithm X

For evaluating the routine `Expected ScaledSFS`(*Q, ℒ, π*^0^, 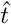, *ω, δ*), which given the data structures associated with the Markov process and an anterior distribution of states *π*^0^, updates an approximate solution 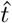 to problem (22) using a SOR(*ω*) iterative scheme until ∥res∥_∞_ *< δ*, and returns que corresponding value of 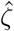.

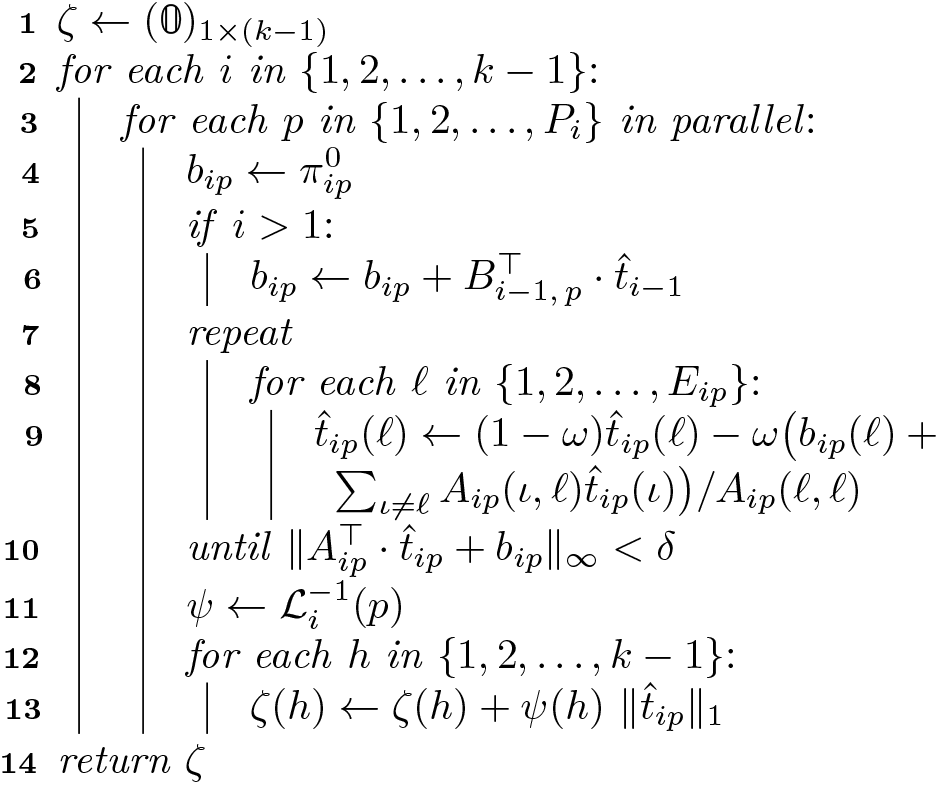

However, the “end user” of Algorithm X is not necessarily interested in 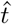 or the residue, but rather in 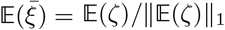, so it would be more meaningful to control the norm of the error 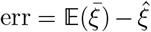, where 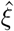 is the approximation of 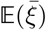 given by Algorithm X: 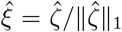. Our goal for this section is therefore to find a way of controlling ∥err∥_∞_ by controlling ∥res ∥_∞_.

A lower bound for ∥err ∥_∞_ can be derived as follows:

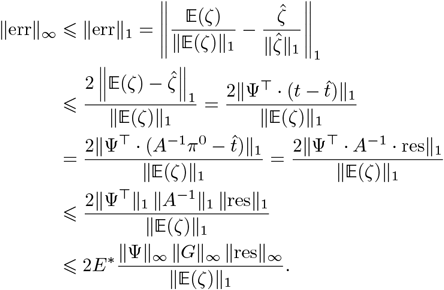

We note that since ‖Ψ‖_∞_ is the largest possible row sum of Ψ, and the rows of Ψ are the contribution vectors of the states of *ε*\*, then ∥Ψ ∥_∞_ = max_*α*∈ *ε*\*_ |*α*| = *k*. We also note that ∥*G* ∥_∞_ = ∥ (*Q*^*^)^−1^∥_∞_. With this we have

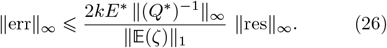

In order to continue, we need an upper bound for 𝔼(*ζ*)_1_. In what follows we use the fact that since Ψ ⩽0 and *G*^⊤^ = *A*^−1^ ⩾0, then all of the coefficients of *G* ·Ψ are also non-negative.

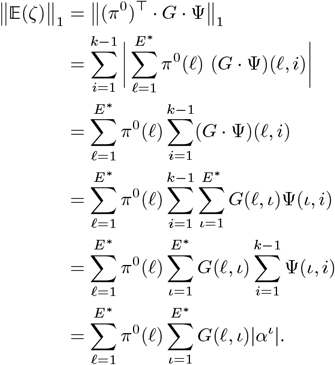

Since *ι ⩽ E*^*^, *α*^*ι*^ is a transient state and therefore has at least two live lineages, thus:

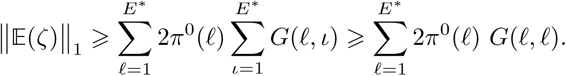

The coefficient *G*(*ℓ, ℓ*) is 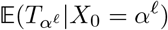, the expected total time the process spends in state *α*^*ℓ*^ prior to absorption, given that the process started in that same state. This expected time can be computed as the product of the total expected number of visits to state 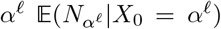, multiplied by the expected duration of each visit, which is an exponentially distributed time with parameter *Q*(*ℓ, ℓ*). An upper bound for *G*(*ℓ, ℓ*) thus can be obtained by observing that 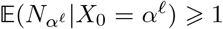, since at least one visit is expected at state *α*^*ℓ*^ in lieu if it being the initial state. Additionally, we note that:

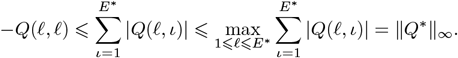

This implies that the expected duration of one visit to state *α*^*ℓ*^ is greater than 1*/* ∥*Q*^*^∥_∞_. With this we have:

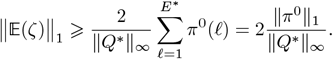

Returning then to (26):

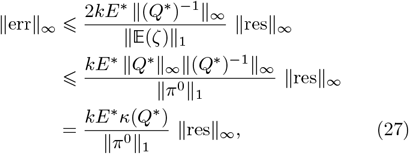

where *κ*(*Q*^*^) is the condition number of the matrix *Q*^*^. These results can be summarized in the following

#### Lemma 1.

*For any *ε* >* 0, *it is sufficient to choose δ as*

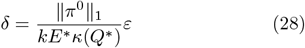

*to have* ∥res ∥_∞_ *< δ* ⇒ ∥err ∥_∞_ *< ε*.

Having a clear criterium for how to control the global error of the normalized expected SFS 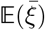, we can proceed to present the main and final algorithm of this section:

#### Algorithm R

For evaluating routine `Expected SFS`(*k, n, ℳ, 𝒮, α*^0^, *ω, ε*), which given the model parameters and an initial sampling state *α*^0^ ∈ *ε*_1_, returns an approximation of 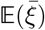 with global error ∥err ∥_∞_ *< ε* using a SOR(*ω*) iterative scheme.

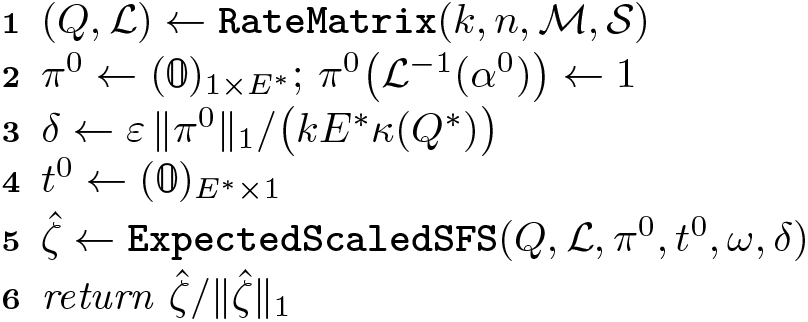

We note that our implementation (SISiFS) does not use the error control mechanisms described in this section, opting instead for the more simple stopping criterion of 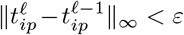. This alternative method avoids the need for the vector-matrix multiplication needed to compute the residue 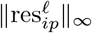, and avoids the need for estimating the condition number *κ*(*Q*^*^). The tradeoff is reduced precision in the error control. These error estimates are showcased in Figure 3.

**Figure 3:**
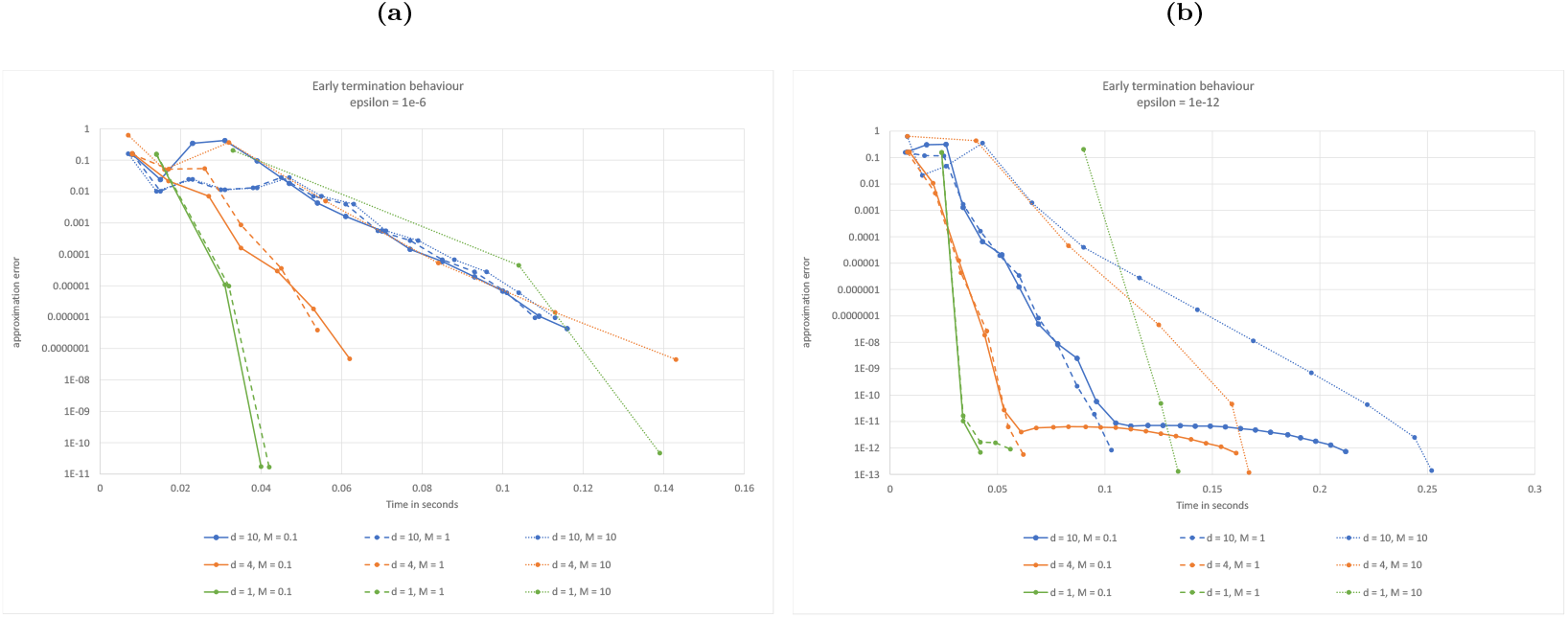
Method error as a function of the computation time. Each line represents one execution of algorithm R’, which iteratively computes the expected SFS of a sample of size *k* = 20 under a symmetrical *n*-island model with *n* = 30. Each highlighted point on the line indicates a partial solution, where the current estimated error is given by the *y*-coordinate, and the time spent so far by the *x*-coordinate. In every case, the algorithm terminates at the first point with a *y*-coordinate lower than *ε*. The solid, dashed and dotted lines correspond to models with a migration rate of *M* = 0.1, 1 and 10 respectively. The green, orange and blue lines correspond to executions of the algorithm with introspection granularity values of *d* = 1, 4 and 10 respectively. Higher values of *d* allow for more intermediate steps (i.e., more opportunities for early termination) at the cost of convergence rate. The value of *ε* is 10^−6^ for panel **(a)** and 10^−12^ for panel **(b)**.

### 3.3. Early termination

The following discussion is concerned with the eventual use of Algorithm R in the context of demographic inference, where early termination is desirable.

Given a value of *ε*, Algorithm R computes a numerical approximation of 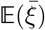 with an error ∥err∥_∞_ *< ε*. However, the internal solution method we use for each of the *P* sub-sub-problems (25) is iterative, meaning that progressively-accurate approximations for which *t*_*ip*_ are being computed, which in turn could be used to compute progressively-accurate approximations of 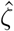. Obtaining early approximations of 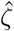 (i.e., approximations where ∥err∥_∞_ is not smaller than *ε yet*) could be useful for implementing more flexible stopping criteria, where we allow the method to converge to the desired high accuracy *ε* only if the SFS is approaching a previously designated *target* SFS; and otherwise terminate the process prematurely, thus saving computational resources.

The key insight for our proposal comes from the observation that Algorithm X operates by simply *updating* or *replacing* a starting solution *t*^0^ with a more accurate one 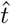, for which ∥res∥_∞_*< δ*. This means that it is possible to continue refining the solution by repeatedly invoking the routine, each time passing as *t*^0^ the final result 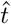 from the previous invocation, and a smaller value of *δ*.

One possible realization of this idea is as follows: suppose we are interested in observing *d* intermediate solutions 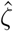 prior to the convergence with ∥err∥ *< ε*, then we may define *d* intermediate tolerances:

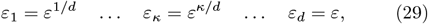

along with the corresponding *δ*_*κ*_ = *δ*^*κ/d*^, and invoke Algorithm X *d* times: once for each of the *δ*_*κ*_. After each invocation we evaluate the corresponding early approximation of 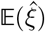, and decide whether to continue or terminate the process. The sequence (29), while certainly not unique, satisfies the desirable properties that the *ε*_*κ*_ are in descending order (assuming *ε <* 1), and are uniformly log-spaced. This implies that the subsequent invocations of Algorithm X, which improve the accuracy of the solution from *δ*_*κ*_ to *δ*_*κ*+1_, should take similar times to execute.

There is a tradeoff to be balanced between execution time and level of control when choosing the value of *d*, since the products 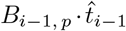 and 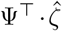 (lines [6] and [12 … 13] of Algorithm X, respectively) have to be computed *d* times each (see Figure 3).

The following revision of Algorithm R takes an additional parameter (*d*), and visits *d* increasingly-accurate approximations of 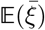.

#### Algorithm R’

Which overloads routine `Expected SFS`(*k, n, ℳ, 𝒮, v, ω, d, ε*) for early termination.

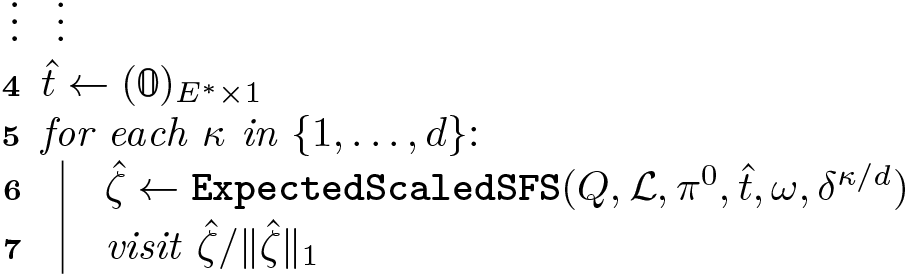

We can see in Figure 3 the tradeoff between computation time and error introspection granularity in an *n*-island model. In general, there are diminishing returns with values of *d* larger than 10, although the exact optimal value depends heavily on the application. In all cases, the fastest results are achieved with the fewer introspection steps.

## 4. Model specialization: Symmetrical *n*-island

Despite the efficient algorithms presented in the previous sections, the state space grows too quickly and becomes computationally intractable even for small values of *k*. For this reason, we believe that the most sensible application for these methods is through *model specializations*. A model specialization is a way of compressing the state space by taking advantage of the symmetries present in a specific demographic model. In this section we use the *n*-island model as a case study for introducing model specializations.

The *n*-island model represents the worst case scenario regarding the size of the state space. Indeed, since all islands are reachable from all other islands, the migration matrix ℳ is completely dense, which in turn implies that the transition rate matrix *Q* is the largest and least sparse possible. On the other hand, the *n*-island model is the most symmetrical demographic model possible: all islands have the same sizes and connectivity, thus they are only distinguishable through the number and configuration of lineages present. Consider for instance the following states in a symmetrical 3-island model with *k* = 4 lineages:

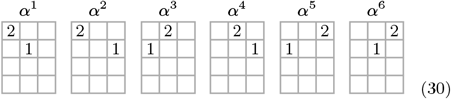

The states in (30) are different only because we have numbered the islands, but since the islands are indistinguishable, they can all be grouped in the same equivalence class. This equivalence class is characterized by one of the islands having two lineages of weight 1 and another island having one lineage of weight 2. This class can be represented by a state matrix where the columns no longer indicate the island number, but are simply an arbitrary indexing of the non-empty islands. We can define a unique class representative by sorting the columns of the equivalent state matrices. We use the reverse-lexicographic ordering on the columns (read from top to bottom). With this ordering, the class representative of the equivalence set (30) would be *α*^1^.

We will assume the existence of a function **ClassRep-resentative** that maps *ε* onto itself, and assigns to every state matrix its class representative. In this section we present modified versions for some of the previously derived algorithms in order for them to work for the *n*-island model specialization.

The results shown in this section were obtained with the software SISiFS (short for Symmetrical Islands Site Frequency Spectrum). The implementation was done using the D programming language, and can be found in the public repository github.com/arredondos/sisifs.

### 4.1. The rate matrix, revisited

We saw in §2.2 that the model state space is discovered by generating all possible migration and coalescence events (algorithms M and C, respectively). In order to adapt these for a symmetrical *n*-island specialization, all we have to do is to only visit the states class-representatives, which as discussed previously, are obtained by sorting the columns of the state matrices. Therefore, the adaptation will consist of, given a destination state *β*, generating its class representative *β*^*^ and visiting it instead.

#### Algorithm sM

For evaluating routine **Symmetrical** `MigrationDestinations`(*m, α*), which given the migration rate *m* and a state matrix *α* of size *k× n*, visits all the tuples (*β, m*) where *β* is a class-representative state satisfying (6) for symmetrical island models, and *m* is the corresponding transition rate.

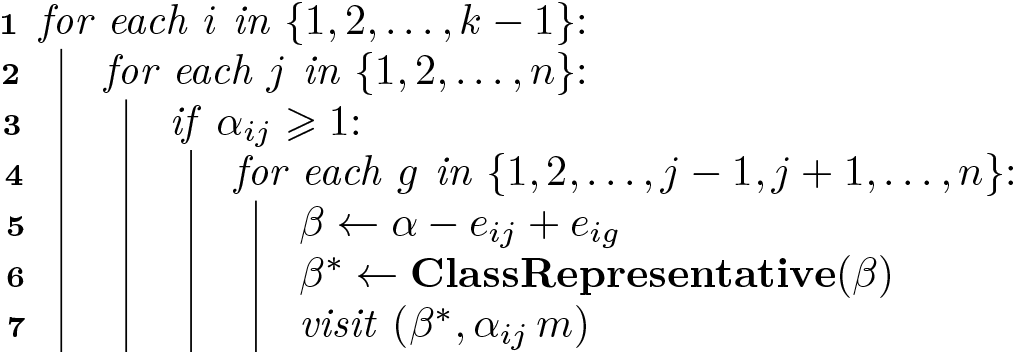

This simple modification allows for a large compression of the state space. See Figure 4 for a comparison of the number of states of the Markov systems associated to the symmetrical *n*-island model, and the most general demographic model possible, where all the states are present.

**Figure 4:**
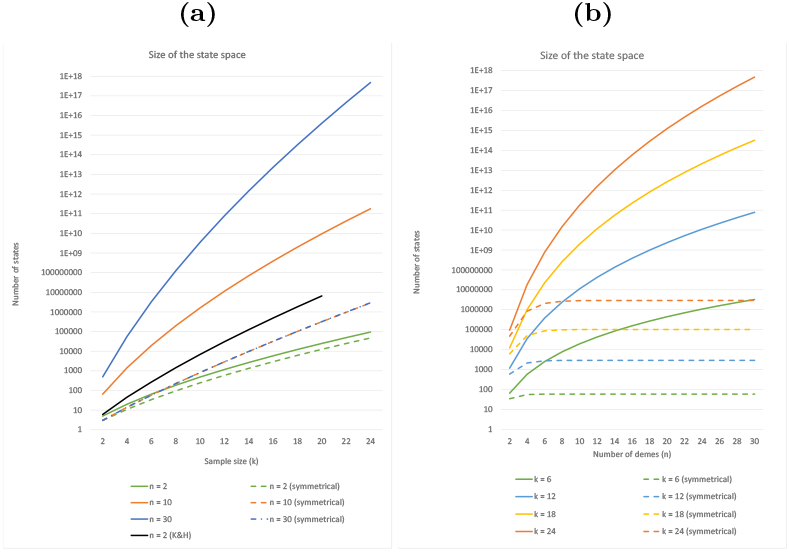
The size of the state space in various Markov processes associated with *n*-island models. Panel **(a)** shows the number of states as a function of the sample size *k*. As a point of reference, we include in this panel the number of states in the isolation-with-migration model of Kern and Hey [19], which computes the exact expected joint SFS of two populations (black plot, data extracted form Figure 2). Panel panel **(b)** shows the same information as a function of the number of islands *n*. We see that with the state space compression (dashed plots), the number of states does not depend on the number of demes once *n k*, as expected.

#### Algorithm sC

Defines `SymmetricalCoalescence Destinations`(*s, α*), which given the deme size *s* of a symmetrical island model and a state matrix *α*, visits all the tuples (*β, c*) where *β* is a class-representative state satisfying (7) for this model, and *c* is the corresponding coalescence rate.

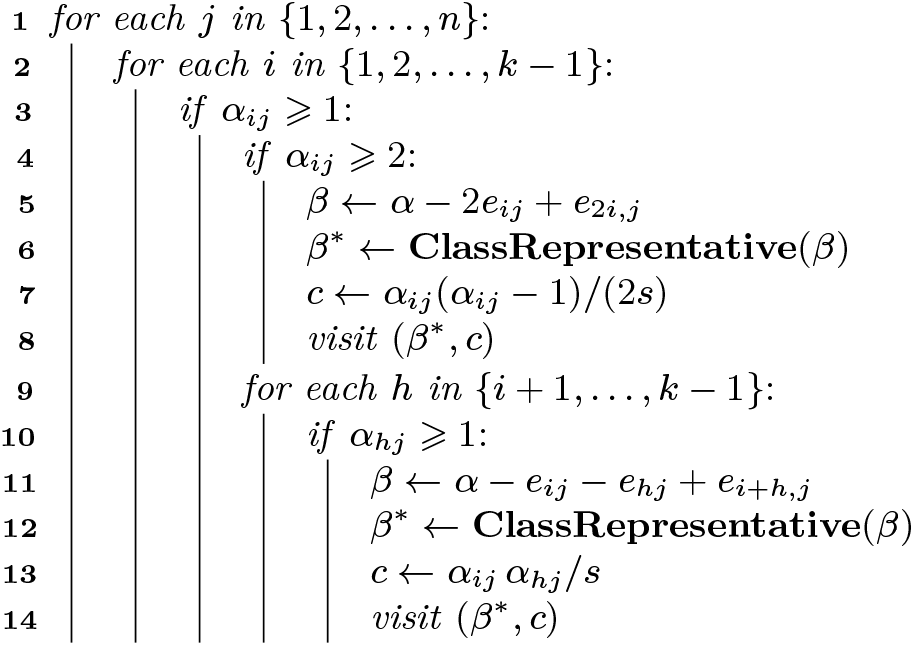

The simplification of the Markov system provided by the *n*-island model specialization goes beyond simply reducing the number of states. Indeed, since now all migration events happen with the same rate *m*, and all coalescence events with the same rate 1*/s*, the coefficients of the rate matrix also become greatly simplified. We can observe in Algorithm sM (line 6) that the coefficients of the migration blocks *A*_*i*_ are of the form ∑_*ℓ,ι*_ *α*_*ℓ,ι*_*m* = *m C*_*ℓ,ι*_. Similarly for the coalescence blocks *B*_*i*_ (see algorithm sC, lines 7 and 12), the transition rates are of the form 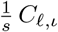. Consequently, we can decompose the rate matrix *Q* of a symmetrical *n*-island model in the following form:

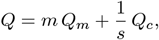

The matrices *Q*_*m*_ and *Q*_*c*_ hold the coefficients *C*_*ℓ,ι*_ for the migration and coalescence blocks, respectively, and they are invariant given *k* and *n* fixed. We can use this representation in order to greatly reduce the computational cost of building the rate matrix (algorithm Q) for the same values of *k* and *n* but many different values of *m* and *s*. See figure 5 for a comparison of the computational resources required to execute algorithm Q and those required to build the rate matrix given *Q*_*m*_ and *Q*_*c*_.

**Figure 5:**
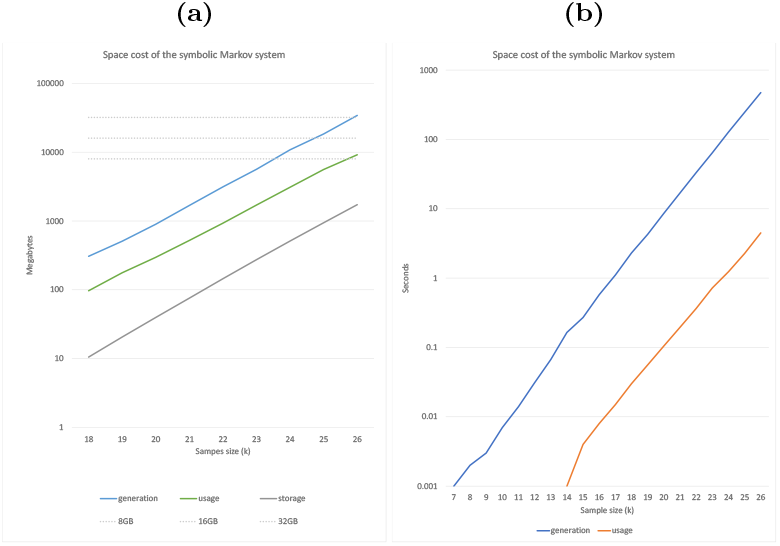
Computational costs associated with the Markov state system for various *n*-island models as a function of the sample size *k*. Panel **(a)** shows the maximum space required in main memory to generate and use the transition rate matrix as per Algorithm Q (blue line). In the case that a symbolic representation of this matrix is available, then the space required in main memory to use it for computing the expected SFS as per Algorithm X is shown as the green line. The grey line shows the space required for storing this symbolic representation of the rate matrix. Panel **(b)** shows the time in seconds needed to generate the rate matrix as per Algorithm Q (blue line) and the time needed to load this matrix from storage (orange line). We note that the space and time costs depicted by the blue lines need only be incurred once for each unique pair of (*k, n*) values.

All time and memory benchmarks presented in this and the next sections were performed on a Ryzen 3700X system with 32GB of RAM. As can be seen in Figure 5, the RAM capacity was the eventual limiting factor in how big a sample could be processed (*k* = 26 in this case). We note that larger samples could be processed by having systems with larger main memory compute the symbolic rate matrix representation remotely and then distribute this file to the local system where the analyses are being conducted. The size of this file, and the RAM cost to load it is also displayed in Figure 5.

### 4.2. The expected SFS in the n-island model

In this section we focus on the problem of computing the expected SFS in the *n*-island model. Specifically, we are interested in validating the correctness of algorithms X and R’, with the *n*-island-specific modifications introduced in §4.1, as well as benchmarking their performance.

We begin by taking a look at the expected SFS itself. Figure 6 shows the plot of 𝔼(*ζ*)*/* ‖𝔼(*ζ*)‖ for various parameter values and sampling patters. Overall, we see that the sampling vector is a key determining factor for the shape of the SFS. Depending on the model parameters, it is relatively common for all or most of the lineages that are sampled in the same deme to coalesce among them before any of them migrates to a different deme. This creates an excess of mutations shared between all the lineages of that deme. This effect can be clearly seen in the figure. For instance, in panel (b) there is a spike at frequency 13 since there are two islands that start with 13 lineages each, and in panel (c) where the sampling vector is [14, 6, 6], there are spikes in the expected SFS at frequencies 14, 6, 14 + 6 and 6 + 6. Another observation is that this effect is amplified when the migration value is small, since it increases the probability that coalescence happens before migration in a deme. Likewise, a smaller number of islands also amplifies this effect of structure, although to a lesser degree than *M*. We can quantify the effect of structure on the SFS by measuring the deviation of the expected SFS in these *n*-island models from the expected SFS under panmixia (1). We present these results in Figure 7. We notice the same spikes near the sampling frequencies, but also that for many of the structure parameters, there is a smaller-than-expected (as compared to a panmictic model) number of unique mutations (singletons and doubletons) and a higher-than-expected number of shared mutations.

**Figure 6:**
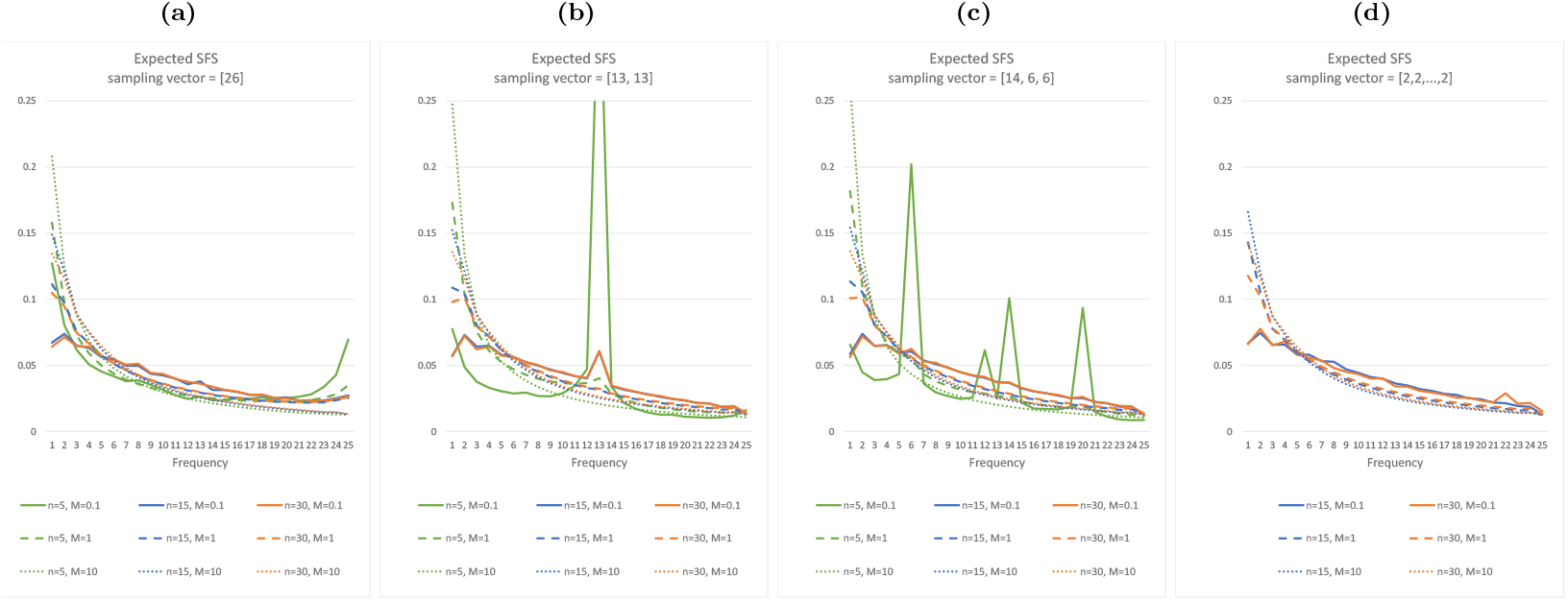
Expected SFS of the *n*-island model for various *n*-island models. All the displayed SFSs correspond to a sample of size *k* = 26 haploids, and the different panels show the effect of different sampling patterns. The curves corresponding to different number of islands are displayed in different colors, and different migration values are displayed in different line styles. We omit the *n* = 5 plots in panel (d) because they are not compatible with the corresponding sampling pattern.

**Figure 7:**
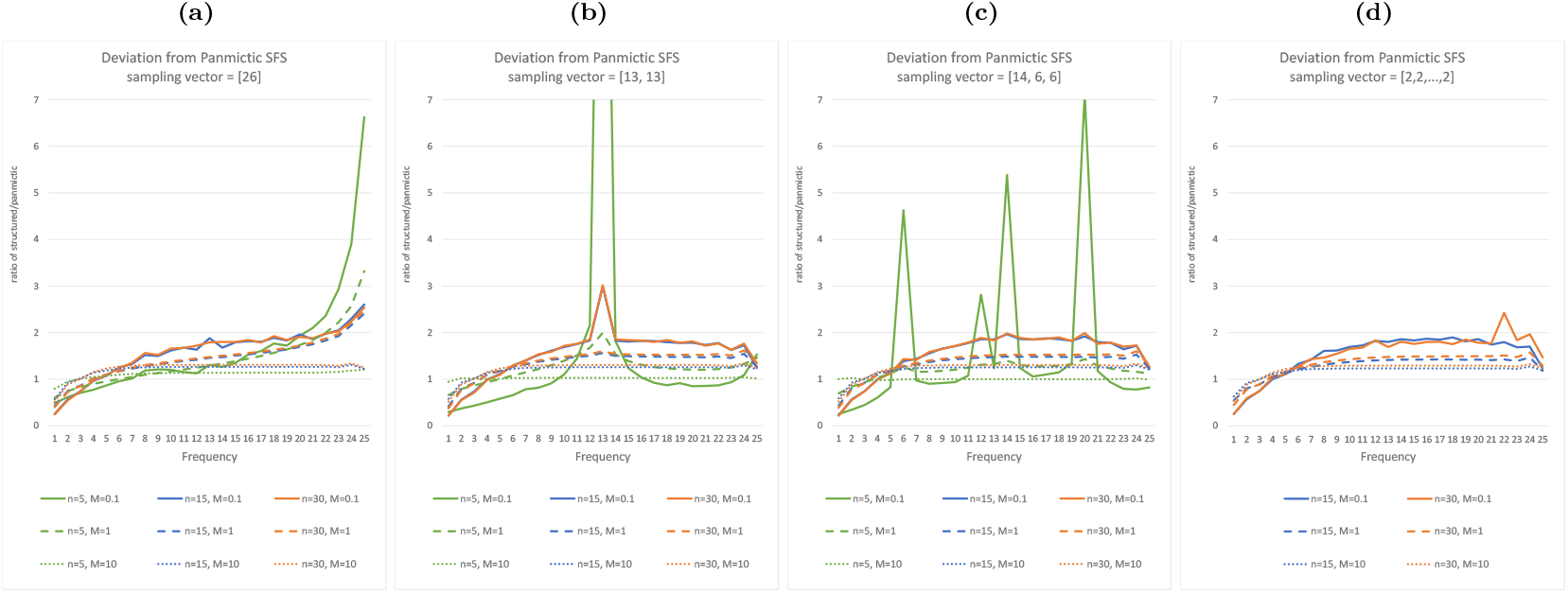
Ratio of the expected SFS in *n*-island models to the expected SFS in panmictic models. Here we show the same *n*-island models and parameters as in Figure 6. See Figure 6 for more details.

These results are not entirely new (see for instance Figure 4.11 in Hein et al. [15]). However, we can now compute these figures exactly (up to machine precision), and very quickly. Figure 8 shows the required time to run Algorithm X for some parameter values of the *n*-island. The sizes of the associated Markov chains can be referenced in Figure 4. For instance, we observe that for a sample of size *k* = 26, the rate matrix has over 8 million states, and the implementation of Algorithm X in SISiFS iteratively solves the associated linear system to a precision of *ε* = 10^−6^ in under 2 seconds, if the migration rate is not too large. The migration rate *M* affects the performance of the method because the condition number of the system matrices *κ*(*A*_*ip*_) increases rapidly with large values of *M*.

**Figure 8:**
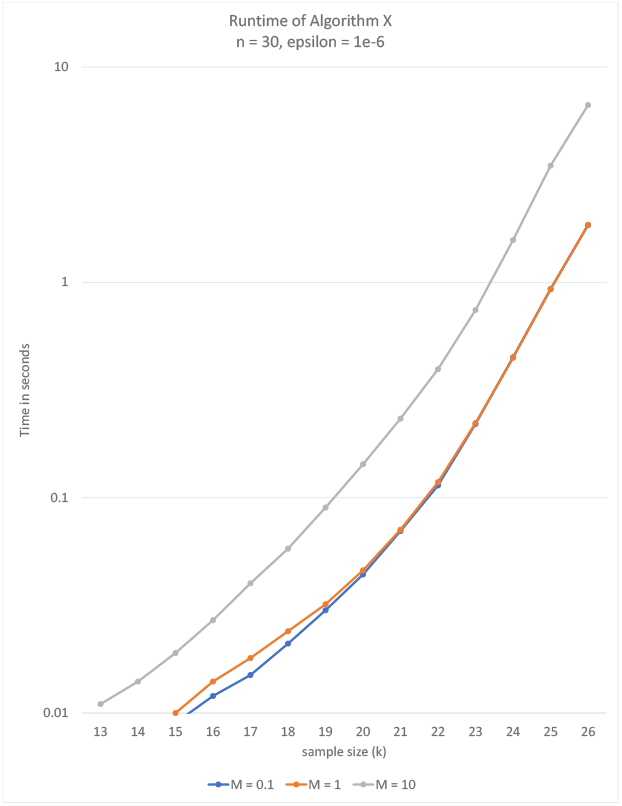
Time to compute the expected SFS in various *n*-island models as a function of the sample size *k*. Each plot shows the time required in seconds to perform a single run of Algorithm X for different values of the migration rate *M*. We note that significantly increasing the value of *M* results in slower convergence. This can be attributed to the increase in the condition number of the rate matrix, which is correlated with the numerical difficulty of the problem. The number of demes is fixed at *n* = 30 so that the size of the state space only depends on the value of *k*, therefore the time complexity can equivalently be read as being a function of the size of the state space. The error tolerance was fixed at *ε* = 10^−6^.

The effect of the high condition number for large *M* can also be observed as numerical instability in Figure 9. This figure showcases the convergence of the expected SFS under various *n*-island models to the expected SFS under panmixia, for which the closed-form expression is known (1) when *M* → ∞. This convergence is expected since island models with a very high migration rate are known to behave like panmictic models with a population size equal to the sum of the island sizes. We verify this fact by making *M* increasingly larger and comparing the two expected SFSs by computing the ∥ · ∥_∞_-based distance between them. We observe that the algorithm converges exponentially to the correct solution in all but the largest cases (*k ⩾*24). This result serves as a validation of Algorithm X, but also indicates that in practice, the degradation of speed and accuracy due to very large values of *M* does not impose a great limitation on the method, given that the expected SFS can be well-approximated by (1) in such cases.

**Figure 9:**
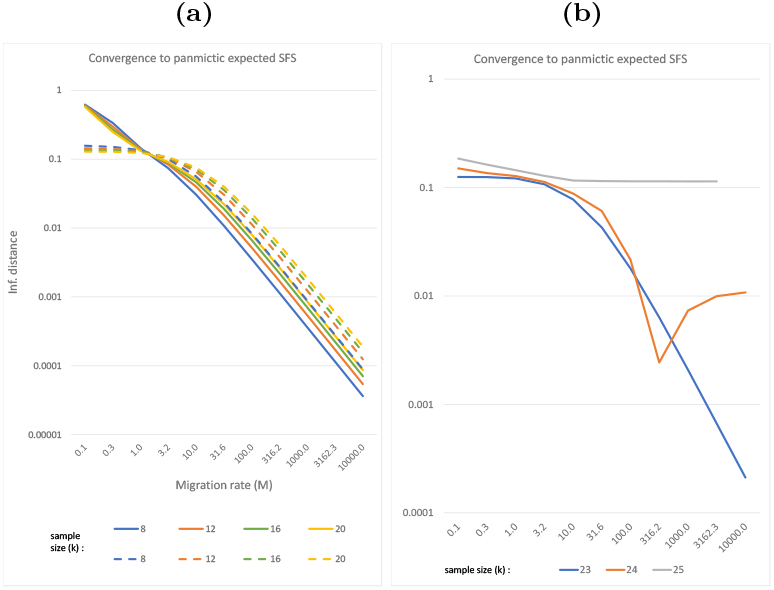
Distance between the expected SFS under panmixia and the expected SFS under various *n*-island models as a function of the migration rate *M*. Panel **(a)** shows the ∥ · ∥_∞_ distance between the expected SFS as computed by Algorithm X and the one expected under panmixia (1) for various sample sizes *k*. The exponential convergence is expected under the presupposition that an *n*-island model with very high migration rate functions equivalently to a panmictic model where the size of the population is the sum of the sizes of the demes. The different plot clustering is due to different sampling schemes: (*k*, 0, …, 0) for the solid plots and (*k/*2, *k/*2, 0, …, 0) for the dashed plots. Panel **(b)** shows the same information for larger sample sizes. We observe that the numerical accuracy of the method degrades for very large values of *M* in these cases. This may be attributed to the increase in the condition number of the rate matrix, which is associated with the numerical difficulty of the problem.

As an additional form of validation, we conducted simulations using the ms software and computed the empirical SFS for samples of various sizes under the *n*-island model. We simulated independent chromosomes continuously with a given value of *θ* until we achieved a certain number of observed segregating sites. After that, all the information at these sites was aggregated and an empirical SFS was computed. The goal is to compare the obtained results with the ones given by our methods. This approach validates both algorithms Q and X. Figure 10 shows the results of these comparisons. We find that estimating the expected SFS with this method is a poor substitute for exactly computing it using our approach for the tested sampling sizes (*k ⩽* 26), both from the numerical accuracy perspective as well as from the required computation time (we note that even though Figure 10 does not show the times required to complete these ms simulations, our method was so many orders of magnitudes faster for the same accuracies, that we felt that a direct comparison was unnecessary).

**Figure 10:**
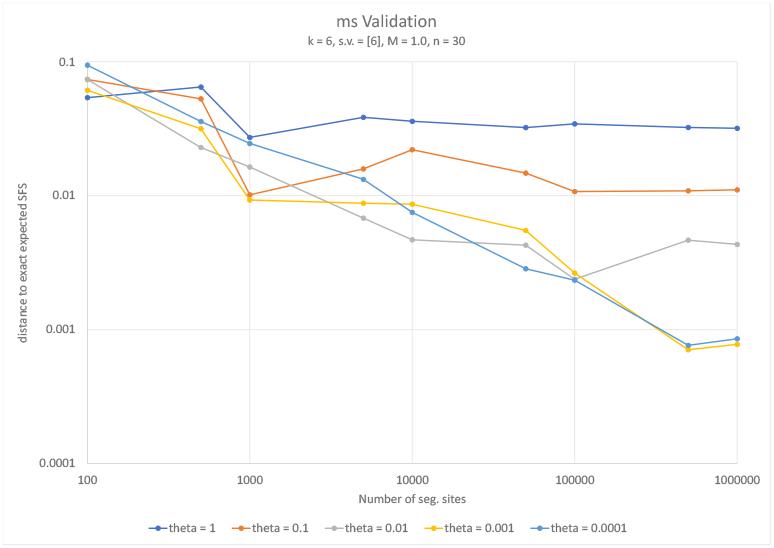
Distance between the expected SFS under one *n*-island model and the empirical SFS obtained form ms simulation as a function of the number of segregating sites. For the representative *n*-island model we chose *n* = 30 islands with a migration rate of *M* = 1 and *k* = 6 lineages sampled in the same deme. The plots show the distance ∥ · ∥_∞_ between the expected SFS for this model and the one obtained by building the observed SFS after many rounds of ms simulations under the same structured model. The different plots correspond to different *θ* values used in the simulations. We notice that lower values of *θ* result in more accurate empirical SFSs as expected, however the simulation approach only achieves acceptable accuracy after a very large number of simulations (note that for lower values of *θ*, a greater number of simulations is required to achieve the same amount of segregating sites).

Figure 3 showcases the early-termination feature of Algorithm R’ (see §3.3). Here, we can see the trade-off between the speed of convergence and the flexibility afforded by the error introspection. Increasing the value of the parameter *d* allows for additional intermediate results of the expected SFS to be available. This could be useful for terminating the computation of the expected SFS if it is not converging towards the desired curve. For instance, in the context of a hypothetical demographic inference application, the computation of the SFS could be aborted before reaching the accuracy specified by *ε* if the expected SFS is not converging towards the target (observed) SFS, and the search algorithm could then focus on testing other parameters.

As expected, the fastest results are achieved with *d* = 1, but since the convergence is often achieved in one or two iterations, it also allows for minimum flexibility. This is the recommended value for one-time computations. In general, the optimum value of the *d* parameter will be heavily dependant on the application that Algorithm R’ is being used in.

## 5. Discussion and future work

The SFS has increasingly become a central and important statistic in demographic inference due to its capacity to succinctly summarize a large amount of information about the genetic diversity of a reasonably large sample in terms of the number of loci and individuals and to the ease with which it can be computed using real data for many species [8, 25]. In this work we have developed a method and several important tools for exactly computing the SFS in the context of a large family of structured demographic models. The proposed method allows for an arbitrary number of demes and an arbitrary connectivity matrix between the demes. We also provide an implementation for the special case of the *n*-island model which we showed was superior to simulation-based estimates of the SFS in both runtime (for the tested sample sizes) and accuracy. It was indeed both surprising and impressive to see that simulation-based approaches tested here had an error threshold which did not appear to decrease when the number of simulations increased (see Figure 10). We also found that simulation-based estimates of the SFS were significantly slower for all tested sample sizes. This suggests that it should be possible to develop an inferential approach similar to that implemented in the SNIF software of Arredondo et al. [1], where the summary statistic would be the SFS rather than the IICR or its estimate, the PSMC of [22]. There are two important questions that would need to be researched here: how to manage the sampling vector and how to choose the distance function.

Several previous studies have exploited the SFS for inferring models with multiple populations. A few of them used similar Markov models as those considered here [30, 4, 19], either assuming a large number of populations without gene flow, or two populations with gene flow. Other studies used different approaches such as simulations or diffusion equations [12, 8, 19]. All these previous studies focused on the joint SFS, where information regarding the origin of the lineages is preserved (or assumed to be known, see below) along with the frequency of the mutations. This approach limits the number of populations that can be modeled, since a high-dimensional joint SFS is both computationally intractable and might become uninformative in practice, since one would expect that most of the coefficients would be null due to lack of information.

In this work we show that by focusing on the aggregate SFS of structured populations, it is possible to have a summary statistic that allows for complex models of population structure and is sensitive to the model parameters (see figures 7 and 6). We also show that by using phase-type theory (see [17]), it is possible to exactly compute this SFS for moderate sample sizes very quickly. We did this by introducing a generalization of Herbots’s structured coalescent that tracks information regarding the ancestry of the lineages as well as their location in the demes. The challenge of the approach is tied to its computational requirements, since the state space of the Markov process grows almost exponentially with the sample size. To combat this effect, we recommend the usage of model specializations. These are Markov processes that are derived from the general process described in Section §2 that include state-reduction optimizations specific to the demographic model in use. These state reductions are achieved by exploiting the symmetries often exhibited in several important classical structured models. We explored the *n*-island model specialization in Section §4. One natural way to continue this line of research is thus to explore other model specializations such as the stepping stone family of models, which would enable us to draw more spatial-aware conclusions from the potential demographic inference applications.

The performance results shown in figures 4 to 3 are limited to the *n*-island model specialization. We do not know yet how these techniques will perform under different models. On the one hand, the *n*-island is the most symmetrical of the structured models, therefore it allows for the greatest compression on the state space, so from this perspective, other models would be expected to require larger state spaces and thus more computational resources or smaller sample sizes. On the other hand, the *n*-island model has the densest migration matrix of all structured models (note that in the stepping stone model for instance, the migration matrix itself is sparse). This means that other models will generate even sparser rate matrices. Given that the computational cost in sparse-algebra methods are associated with the number of non-null coefficients, this could counter the effect of the increased state space. It is hard to estimate how these two opposite effects will ultimately affect the performance of the methods. This is why more specializations have to be developed and tested.

An important but neglected issue in demographic inference is how inference can be influenced by the sampling scheme [29, 7, 23]. This has consequences on how the sampling vector is defined (and assumed to be) during the inferential process. This sampling vector is important for any structured demographic model that incorporates or rather defines multiple samples. We have seen here that it can have a profound impact on the SFS of otherwise identical *n*-island models (indeed, some of the features in Figure 6 could be wrongly interpreted as the effect of selection). Such is the case as well for the IICR_*k*_ (see the differences between IICR_*s*_ and IICR_*d*_ in Chikhi et al. [6]). We stress that it is also an issue for studies that use the isolation with migration model for various reasons. First the use of an IM model implicitly assumes that we know from which populations the samples were obtained, when in reality samples may be aggregated to obtain large enough sample sizes for the different “populations”. Second, it is assumed that only the populations sampled and modelled are important and that other non sampled populations will not significantly influence demographic inference. This stresses that the demographic model and the sampling vector are not independent and that any demographic inference process requires decisions that may have significant consequences on the history that we first reconstruct and later tell [7, 23, 6, 5].

One initial approach is to assume that the sampling vector is known, although this defeats the purpose of a general inferential approach. Alternatively, we could attempt to infer the sampling vector. This would add a great burden of dimensionality to the parameter space which could render the method impractical. Perhaps a compromise can be achieved between these two extreme approaches, where any existing information about the samples is incorporated in the model (for instance, lineages 1 and 2 are always from the same but possibly unknown deme, as are lineages 3 and 4, etc.). In models that are spatially aware, such as the many variants of the stepping stones model, an alternative approach to managing the sampling vector is possible. We can assume that the actual (physical) sampling locations are known, given a map of the sampling region. We can then associate each sample with the nearest model deme corresponding to that region of the map, while the spatial model, with varying number of demes would itself be inferred in an iterative process. This approach would allow the sampling vector to be constructed organically from the rest of the model parameters. For instance, as the number of demes would decrease the number of samples integrated to the demes would increase and the model parameters would be thus updated. See Figure 11 for an example. We are not aware that any such method exists today but the joint use of the SFS and of the IICRs of individuals sampled through space might make that inference possible. Indeed, [6] have shown that the IICR from individuals sampled in different demes from a stepping-stone differ in predicted ways as a function of the number of demes, the migration rates, their pairwise distances in the stepping stone and their location in relation to the center and edges. Using jointly the IICRs of individuals sampled in space and the corresponding SFS might thus offer an efficient inferential framework because both statistics can be computed quickly thanks to the methods presented here and in our previous studies [26, 1]. Regarding the question of what distance to use for inference, this will depend on the exact nature of the method. For purely SFS based methods, log-likelihood metrics have been successfully used in the past [12, 8] and could thus be used. Altogether, the developments presented here should be seen as a tiny but first step towards incorporating the SFS as a summary statistic in inference applications for demographic models of structure in which the number of islands is unknown and potentially large.

**Figure 11:**
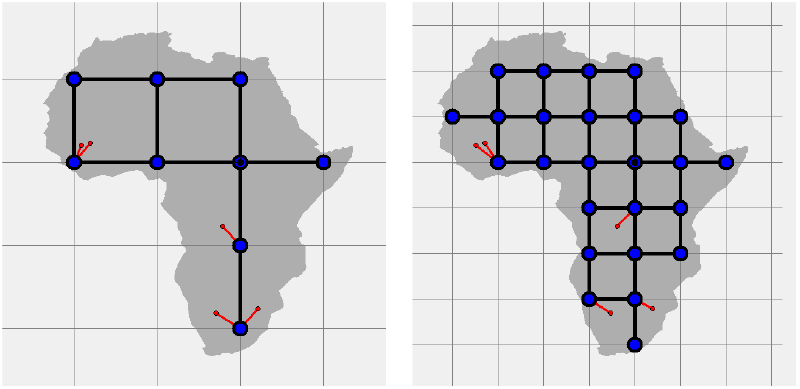
A method for parameterizing spatial structured models. A stepping-stone model is defined by overlaying a grid onto the map of the region of interest. The physical locations of the samples (the red dots in the figure) are assumed to be known, and each one is associated with the nearest deme (the association is represented by a red line connecting the sample with a deme). In this example model, it is possible to specify the number of islands, their connections, and the sampling vector by setting a single parameter: the grid spacing.

## References

[1] Arredondo, A., Mourato, B., Nguyen, K., Boitard, S., Rodríguez, W., Noûs, C., Mazet, O., Chikhi, L., 2021. Inferring number of populations and changes in connectivity under the n-island model. Heredity 126, 896–912.

[2] Beaumont, M.A., 1999. Detecting population expansion and decline using microsatellites. Genetics 153, 2013–2029.

[3] Bladt, M., Nielsen, B.F., 2017. Matrix-exponential distributions in applied probability. volume 81. Springer.

[4] Chen, H., 2012. The joint allele frequency spectrum of multiple populations: a coalescent theory approach. Theoretical population biology 81, 179–195.

[5] Chikhi, L., 2023. Basato su una storia vera: come fanno i genetisti delle popolazioni umane a conoscere le storie che raccontano?, Accademia Nazionale dei Lincei, Roma. pp. 181–211.

[6] Chikhi, L., Rodríguez, W., Grusea, S., Santos, P., Boitard, S., Mazet, O., 2018. The IICR (inverse instantaneous coalescence rate) as a summary of genomic diversity: insights into demographic inference and model choice. Heredity 120, 13–24.

[7] Chikhi, L., Sousa, V.C., Luisi, P., Goossens, B., Beaumont, M.A., 2010. The confounding effects of population structure, genetic diversity and the sampling scheme on the detection and quantification of population size changes. Genetics 186, 983–995.

[8] Excoffier, L., Dupanloup, I., Huerta-Sánchez, E., Sousa, V.C., Foll, M., 2013. Robust demographic inference from genomic and snp data. PLoS Genetics 9, e1003905.

[9] Fay, J.C., Wu, C.I., 2000. Hitchhiking under positive darwinian selection. Genetics 155, 1405–1413.

[10] Fu, Y.X., 1995. Statistical properties of segregating sites. Theoretical population biology 48, 172–197.

[11] Griffiths, R.C., Tavaré, S., 1998. The age of a mutation in a general coalescent tree. Stochastic Models 14, 273–295.

[12] Gutenkunst, R.N., Hernandez, R.D., Williamson, S.H., Bustamante, C.D., 2009. Inferring the joint demographic history of multiple populations from multidimensional SNP frequency data. PLoS Genetics 5, e1000695.

[13] Hackbusch, W., 1994. Iterative solution of large sparse systems of equations. volume 95. Springer.

[14] Hardy, G.H., Ramanujan, S., 1918. Asymptotic formulæ in combinatory analysis. Proceedings of the London Mathematical Society 2, 75–115.

[15] Hein, J., Schierup, M., Wiuf, C., 2004. Gene genealogies, variation and evolution: a primer in coalescent theory. Oxford University Press, USA.

[16] Herbots, H.M.J.D., 1994. Stochastic models in population genetics: genealogy and genetic differentiation in structured populations. Ph.D. thesis.

[17] Hobolth, A., Siri-Jegousse, A., Bladt, M., 2019. Phase-type distributions in population genetics. Theoretical Population Biology 127, 16–32.

[18] Hudson, R.R., 2015. A new proof of the expected frequency spectrum under the standard neutral model. Plos one 10, e0118087.

[19] Kern, A.D., Hey, J., 2017. Exact calculation of the joint allele frequency spectrum for isolation with migration models. Genetics 207, 241–253.

[20] Knuth, D.E., 2005. The Art of Computer Programming, Volume 4, Fascicle 3: Generating All Combinations and Partitions. Addison-Wesley Professional.

[21] Lehmann, D.J., Smyth, M.B., 1981. Algebraic specification of data types: A synthetic approach. Mathematical systems theory 14, 97–139.

[22] Li, H., Durbin, R., 2011. Inference of human population history from individual whole-genome sequences. Nature 475, 493–496.

[23] Mazet, O., Rodríguez, W., Grusea, S., Boitard, S., Chikhi, L., 2016. On the importance of being structured: instantaneous coalescence rates and human evolution—lessons for ancestral population size inference? Heredity 116, 362.

[24] Nielsen, R., 2000. Estimation of population parameters and recombination rates from single nucleotide polymorphisms. Genetics 154, 931–942.

[25] Poelstra, J.W., Salmona, J., Tiley, G.P., Schüßler, D., Blanco, M.B., Andriambeloson, J.B., Bouchez, O., Campbell, C.R., Etter, P.D., Hohenlohe, P.A., et al., 2021. Cryptic patterns of speciation in cryptic primates: microendemic mouse lemurs and the multispecies coalescent. Systematic Biology 70, 203–218.

[26] Rodríguez, W., Mazet, O., Grusea, S., Arredondo, A., Corujo, J.M., Boitard, S., Chikhi, L., 2018. The IICR and the non-stationary structured coalescent: towards demographic inference with arbitrary changes in population structure. Heredity 121, 663–678.

[27] Tajima, F., 1983. Evolutionary relationship of dna sequences in finite populations. Genetics 105, 437–460.

[28] Tajima, F., 1989. Statistical method for testing the neutral mutation hypothesis by DNA polymorphism. Genetics 123, 585–595.

[29] Wakeley, J., 1999. Nonequilibrium migration in human history. Genetics 153, 1863–1871.

[30] Wakeley, J., Hey, J., 1997. Estimating ancestral population parameters. Genetics 145, 847–855.

[31] Watterson, G., 1975. On the number of segregating sites in genetical models without recombination. Theoretical population biology 7, 256–276.

[32] Wilkinson-Herbots, H.M., 1998. Genealogy and subpopulation differentiation under various models of population structure. Journal of Mathematical Biology 37, 535–585.

[33] Wright, S., 1943. Isolation by distance. Genetics 28, 114.

